# Pleiotropy promotes the evolution of inducible immune responses in a model of host-pathogen coevolution

**DOI:** 10.1101/2022.08.02.502467

**Authors:** Reese A. Martin, Ann T. Tate

## Abstract

Components of immune systems face significant selective pressure to efficiently use organismal resources, mitigate infection, and resist parasitic manipulation. A theoretically optimal immune defense balances investment in constitutive and inducible immune components depending on the kinds of parasites encountered, but genetic and dynamic constraints can force deviation away from theoretical optima. One such potential constraint is pleiotropy, the phenomenon where a single gene affects multiple phenotypes. Although pleiotropy can prevent or dramatically slow adaptive evolution, it is prevalent in the signaling networks that compose metazoan immune systems. We hypothesized that pleiotropy is maintained in immune signaling networks despite slowed adaptive evolution because it provides some other advantage, such as forcing network evolution to compensate in ways that increase host fitness during infection. To study the effects of pleiotropy on the evolution of immune signaling networks, we used an agent-based modeling approach to evolve a population of host immune systems infected by simultaneously co-evolving parasites. Four kinds of pleiotropic restrictions on evolvability were incorporated into the networks, and their evolutionary outcomes were compared to, and competed against, non-pleiotropic networks. As the networks evolved, we tracked several metrics of immune network complexity, relative investment in inducible and constitutive defenses, and features associated with the winners and losers of competitive simulations. Our results suggest non-pleiotropic networks evolve to deploy highly constitutive immune responses regardless of parasite prevalence, but some implementations of pleiotropy favor the evolution of highly inducible immunity. These inducible pleiotropic networks are no less fit than non-pleiotropic networks and can out-compete non-pleiotropic networks in competitive simulations. These provide a theoretical explanation for the prevalence of pleiotropic genes in immune systems and highlight a mechanism that could facilitate the evolution of inducible immune responses.

**Author Summary:** Genes involved in immune defense are hotspots of adaptive evolution as they resist rapidly evolving parasites and pathogens. Pleiotropic genes, which affect multiple discrete traits, have been shown to evolve at a much slower rate than non-pleiotropic genes but are highly represented in the immune system. The evolutionary effects of pleiotropic signaling genes on immune evolution are poorly understood, so we developed a model of pleiotropic signaling network evolution to address this gap in knowledge. Our results show that pleiotropy may be an important genomic feature in the development of inducible immunity.

## Introduction

Organismal immune systems have been honed over evolutionary time to balance the benefits of infection resistance against the energetic and pathological costs of mounting and maintaining an immune response (1). Optimal investment in immunity varies based on organismal life histories (2–4), but often features constitutive immune activity (5,6), wherein a defense mechanism is always active even when no threat is present. In humans, such quiescent immune activity can exceed 15% of basal metabolic demands, making the immune system one of the most energetically costly body systems, even in the absence of pathogens (7). In contrast to constitutive responses, inducible immune responses are inactive until a threat is detected, at which point they quickly come online, clear the threat, and undergo rapid suppression by negative regulators (8). As such, inducible immunity is a smaller drain on host fitness than constitutive immunity, but the time needed to activate the response may cede the temporal advantage against rapidly replicating pathogens (6,9).

Previous studies attempting to define optimal investment in constitutive and inducible immune responses have suggested that high parasite virulence and predictability, including frequent exposure and fixed within-host growth rates (10), favor the evolution of constitutive immunity (6). Inducible immunity is favored when infection rates are low, parasite growth rates are varied (10), immune effector proteins can be produced rapidly (6), and when parasites are not co-evolving with the host (11). A recent study on the evolution of immune signaling networks suggested that selection may favor constitutive defenses even when an inducible response is theoretically optimal because inducible architectures and their evolutionary intermediates may be less robust to parasite manipulation, reflecting a more general limitation associated with the evolution of network complexity (11). Thus, an open question surrounding the evolution of immune systems is whether there are other factors that could ease the constraints associated with the evolution and robustness of inducible immune responses.

A ubiquitous but puzzling property of many immune signaling networks may provide an answer to this problem or could represent yet another constraint on the evolution of inducibility. Genetic pleiotropy, the phenomenon in which a single gene affects multiple distinct phenotypes, has been described in immune signaling networks from a variety of taxa. In plants, for example, DELLA proteins regulate network crosstalk between gibberellin-mediated growth and jasmonic-acid mediated defense, affecting two traits that compete for limited energetic resources (12). In insects, the Toll pathway is crucial to both developmental patterning in fly embryos and defense against bacterial and fungal infections (13). Recent work has underscored the high frequency of pleiotropy in insect immune and developmental signaling pathways (4,14) and a study of the human genome identified an overabundance of pleiotropy in the immune system (15). Thus, pleiotropy appears to be a common property of immune signaling networks, but it is not clear whether and when it might constrain or benefit adaptive evolution.

Pleiotropy could exacerbate antagonism among traits at both within-host and evolutionary scales, and many empirical lines of evidence emphasize the potential for constraint. For example, the plasticity of pleiotropic gene expression in response to environmental perturbations is restricted relative to non-pleiotropic counterparts (16). Pleiotropy can also impede the efficacy of selection, as purifying selection to retain one phenotype can interfere with substitutions that might provide an adaptive benefit for another (14,17,18); this effect is magnified between traits that contribute equally to organismal fitness (19). We further hypothesize that the elevated rate of purifying selection on some pleiotropic genes (14,20) and their abundance in the immune system could limit immune network adaptability and present stable avenues of manipulation to parasites.

Pleiotropy is always deleterious, however, as computational (21) and genomics studies (22,23) of trait evolution demonstrate that pleiotropy commonly arises as a neutral process associated with the generation of new traits. Pleiotropic compression of genomes could also be vital for genome size management, which has implications for organismal fitness (18,24). While these aspects of pleiotropy positively affect organismal fitness, it remains unclear what other beneficial changes to network evolution may accompany the incorporation of pleiotropic signaling proteins. To address this open question, we constructed an agent-based model of host parasite co-evolution featuring immune signaling networks and parasites that manipulate host signaling.

Our networks include three fundamental components: detectors, signaling proteins, and effectors. In an immunological context, detectors are pattern recognition receptor proteins like peptidoglycan recognition proteins and Toll-like receptors that directly sense parasites and initiate a response. Signaling proteins can be part of extracellular or intracellular signaling cascades that relay information from detector proteins to effector proteins. Effector proteins (granzymes, antimicrobial peptides, etc.) act to kill parasites or inhibit growth (25). We represent the proteins and protein-protein interactions that compose a signaling network as a collection of nodes and edges respectively. The pleiotropic protein explicitly contributes to immune signaling but also implicitly regulates a non-specific trait like development (26,27), such that its evolution is inherently constrained. Here we present four complementary implementations of this pleiotropic protein: 1) random connections between the pleiotropic protein and other proteins in the network that are fixed across evolutionary time; 2&3) a single connection from the pleiotropic protein that up-or downregulates the effector as a side effect of its function in development; and 4) the protein is allowed to evolve but under reduced mutation rates relative to other network components, capturing reduced evolutionary rates associated with purifying selection (Fig. 1).

**Figure 1:**
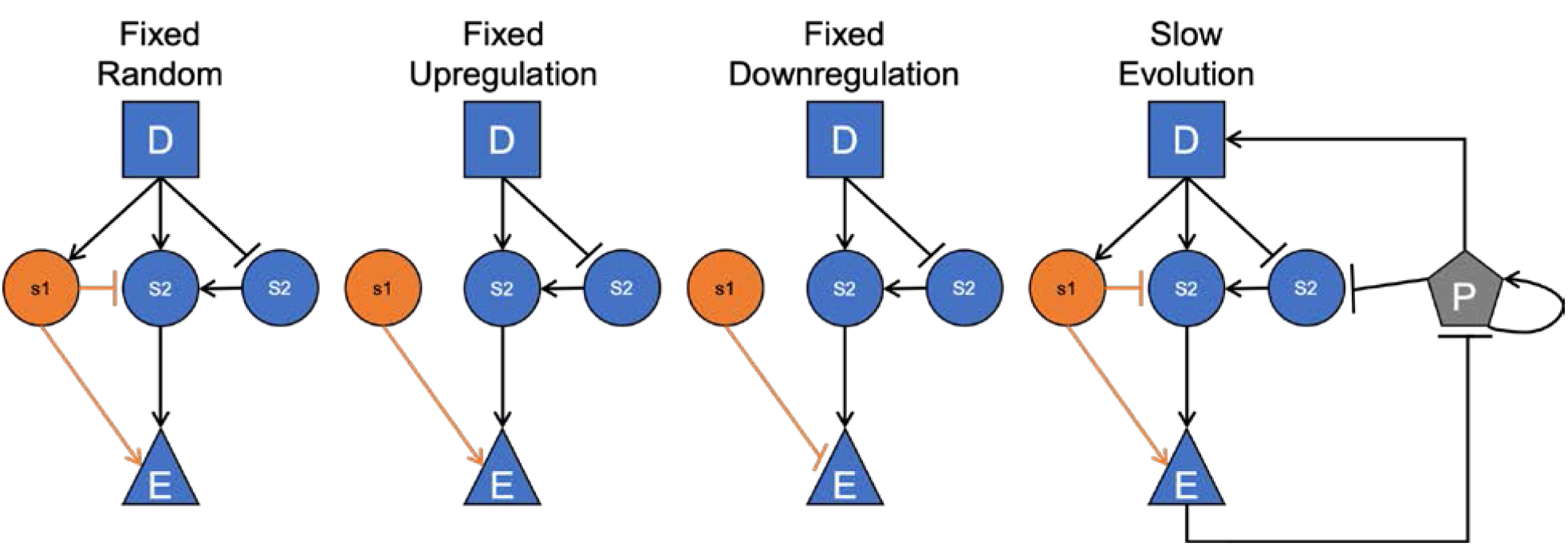
Examples of initial networks, with pleiotropic restraints highlighted. A parasite is shown infecting the Slow Evolution network, where is self-replicates, activates the detector, and is destroyed by the effectors. D = detector, S = signaling protein, E = effector, P = Parasite. Orange arrows show connections fixed via pleiotropic action.

We first investigated whether these implementations of pleiotropy favor distinct patterns of constitutive and inducible immunity over evolutionary time relative to non-pleiotropic networks, keeping in mind that the predictability of infection could influence the outcome. To understand why pleiotropy might be such a common property of immune systems, we then investigated whether pleiotropic networks are capable of outcompeting non-pleiotropic signaling networks over evolutionary time and defined the network properties associated with successful networks. Our results provide a new framework for understanding the evolutionary maintenance of pleiotropy in immune systems and could be applied broadly to understand the evolution of immune signaling networks or the evolution of pleiotropic network architecture among the myriad traits that influence organismal fitness.

## Results

Our agent-based model of signaling network co-evolution features hosts, defined by their immune networks, and parasites that disrupt host signaling to improve their own reproductive success. We present data from 100 simulations for each form of pleiotropic constraint and a non-pleiotropic case at three levels of lifetime infection risk [10%, 50%, 90%], with all simulations consisting of 500 generations. The initial host population was created with random connections between signaling proteins, and each host had the same implementation of pleiotropy within a scenario. Host fitness was evaluated based on immune effector levels pre- and post-infection, parasite burden, and network size (see Materials and Methods equation 2). Parasite fitness was determined by the cumulative magnitude of infection over time within each host. Host evolution allowed for the duplication and deletion of signaling proteins as well as adding, removing, or altering connections between any two proteins in the network. Pleiotropic connections were immutable in most cases, and no direct connection between the detector and effector was allowed. Parasites were allowed to alter the identity and regulatory activity of their targeted signaling protein over evolutionary time.

To evaluate the degree to which hosts could successfully mount an immune defense during infection (network robustness) we compared a time course of effector levels from an intact network to the effector levels during infection when a single signaling protein was removed from the network. To compare the dynamics of the intact and knockout networks, we calculated the mean absolute difference between the two time-courses. To evaluate fundamental properties associated with these networks we measured network nodes, their total connectivity (the number of edges in the network divided by the total possible number of edges), and the distinct paths from the detector to effector protein, where a distinct path does not include a signaling protein used in any other path.

### A higher infection risk favors the evolution of stronger and inducible immune signaling

To determine how pleiotropic proteins may alter immune signaling network evolution, we first needed a baseline understanding of how non-pleiotropic networks evolve. We modeled the co-evolution of non-pleiotropic signaling networks at three chances of infection to uncover the relationship between parasite prevalence and host immune evolution. When the individual chance of infection in a simulation was 10%, most hosts evolved predominantly constitutive immune responses (Fig. 2A) with minimal immune effector activity (Fig. 2B). As the chance of infection increased, the likelihood of hosts developing a mixed-strategy immune response also increased, though predominantly inducible immune responses were still rare. Peak immune effector activity in constitutive responses increased dramatically when the chance of infection increased from 10% to 50% but remained similar in the transition from a 50% chance of infection to a 90% chance of infection. When inducible responses evolved, they tended to show higher peak immune effector activity than constitutive responses (Fig. 2B). The 10% chance of infection produced hosts that were robust to parasitic interference, with a negligible difference in immune effector dynamics between knockouts and intact networks during infection. As the chance of infection increased, we observed a larger discrepancy between intact and knockout networks (Fig. 3). Network size and the distinct paths from the detector to the effector increased with the chance of infection, but network connectivity stayed consistent across all chances of infection (Supplemental Fig. 4-6). Finally, inducibility was not favored when the chance of infection was low, with winners of competitions being no more inducible than losers (Fig. 4).

**Figure 2:**
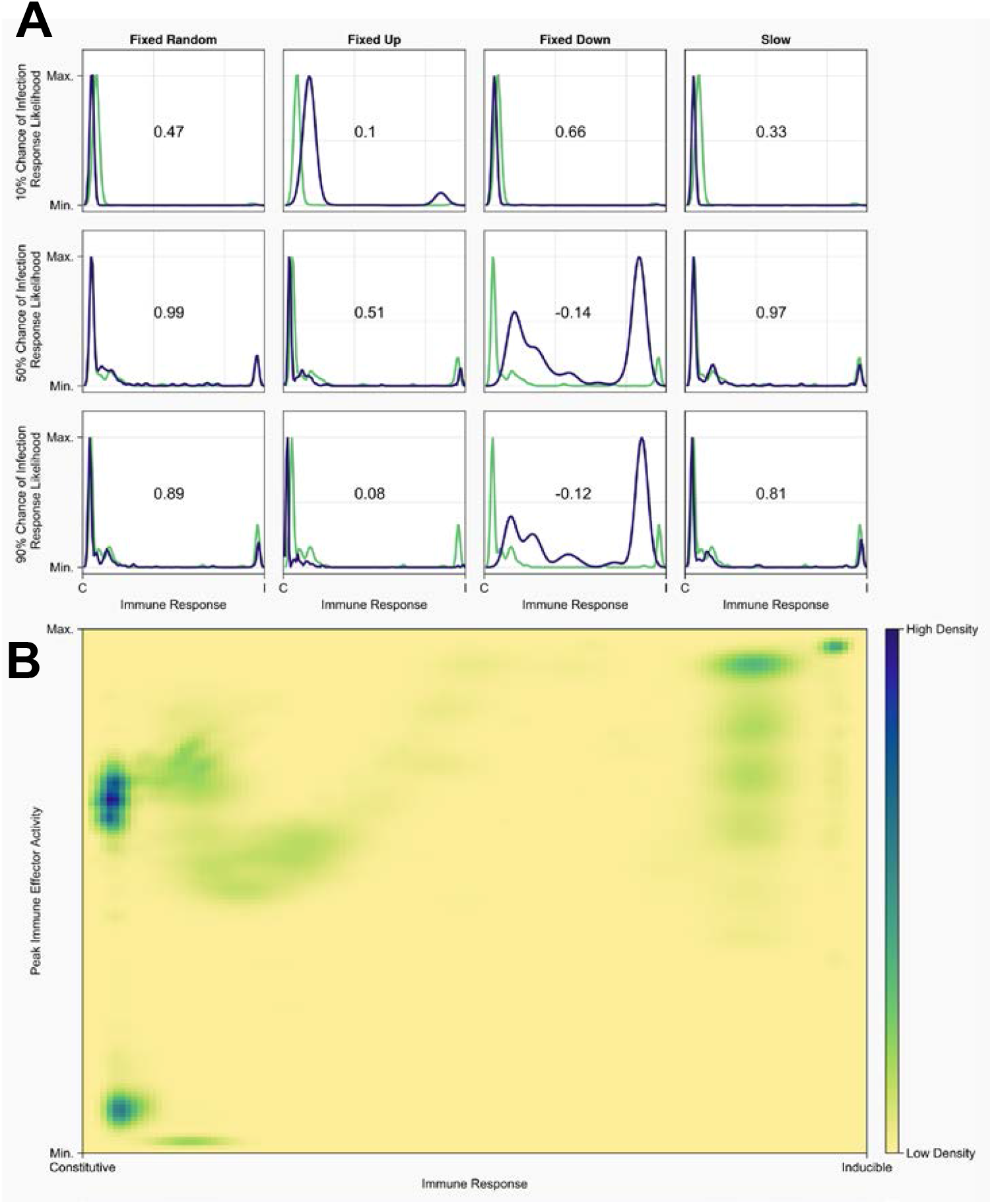
**A)**Normalized probability density function of the proportion of agent immune responses that are induced by parasites. Non-pleiotropic networks are represented in green and pleiotropic networks in blue. All plots on the same row have the same chance of infection (10%, 50%, or 90% descending), and plots in the same column compare the non-pleiotropic network against the indicated pleiotropic constraint. Presented in each plot is the Pearson correlation coefficient calculated between the non-pleiotropic and pleiotropic networks. **B)** Heatmap of the magnitude of maximum immune response attained during infection vs proportion of immune response that is induced by parasites. Darker colors indicate more individuals expressing the magnitude and response combination.

**Figure 3:**
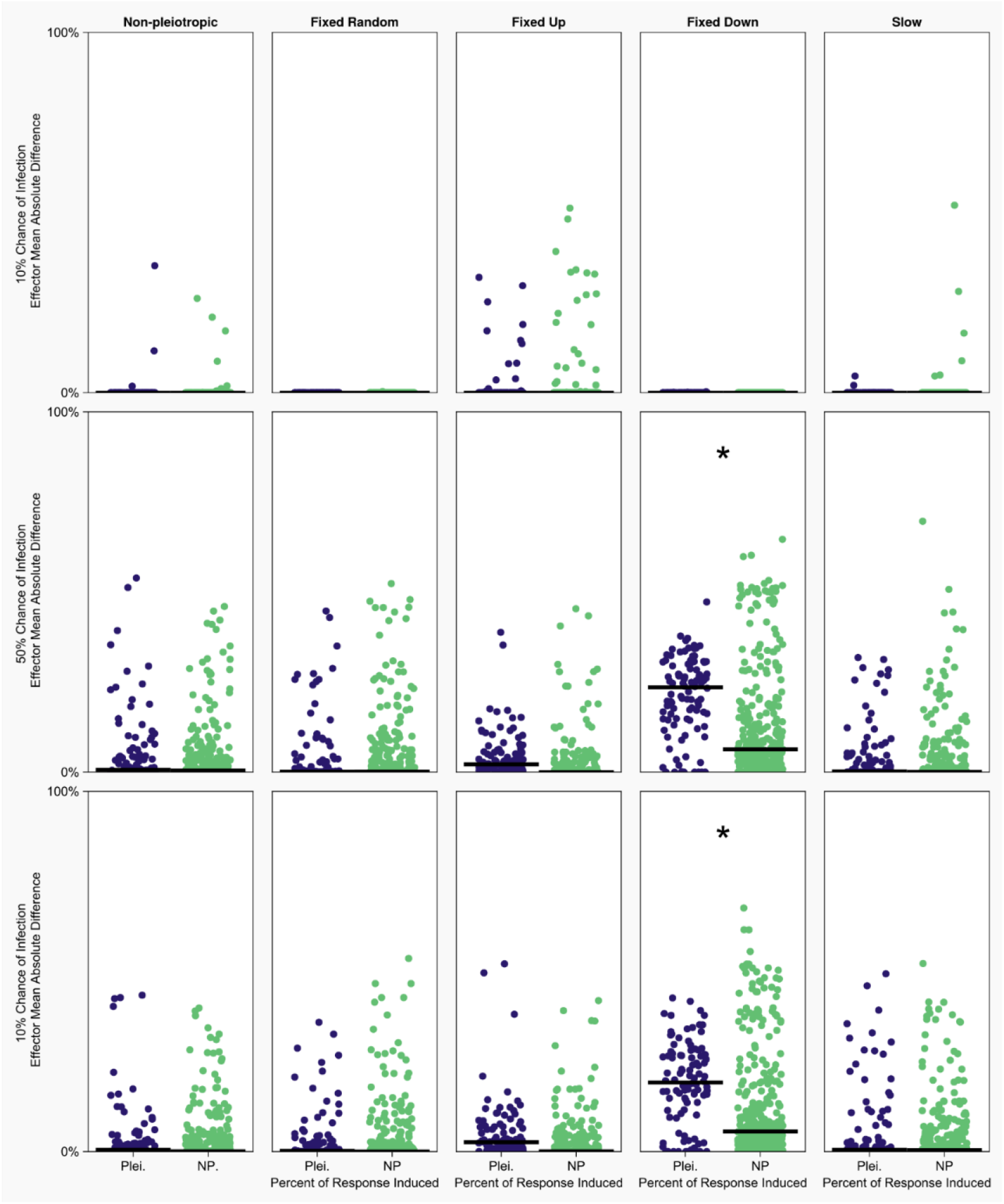
Analysis of network robustness to silenced signaling proteins, measured as the mean absolute difference in effector levels between intact networks and single signaling protein knockout networks. Effector levels were measured during infection by a non-disrupting “dummy” parasite. Blue indicates knockout of the pleiotropic signaling protein, green indicates knockout of a single non-pleiotropic protein. All plots on the same row have the same chance of infection (10%, 50%, or 90% descending), and plots in the same column have the same implementation of pleiotropy. There are no pleiotropic nodes in non-pleiotropic networks (leftmost column) so nodes were just chosen at random twice. The networks used in this analysis were the most common network at the end of the simulation from which they originated. Asterisks denote significant differences between pleiotropic and non-pleiotropic knock outs.

**Figure 4:**
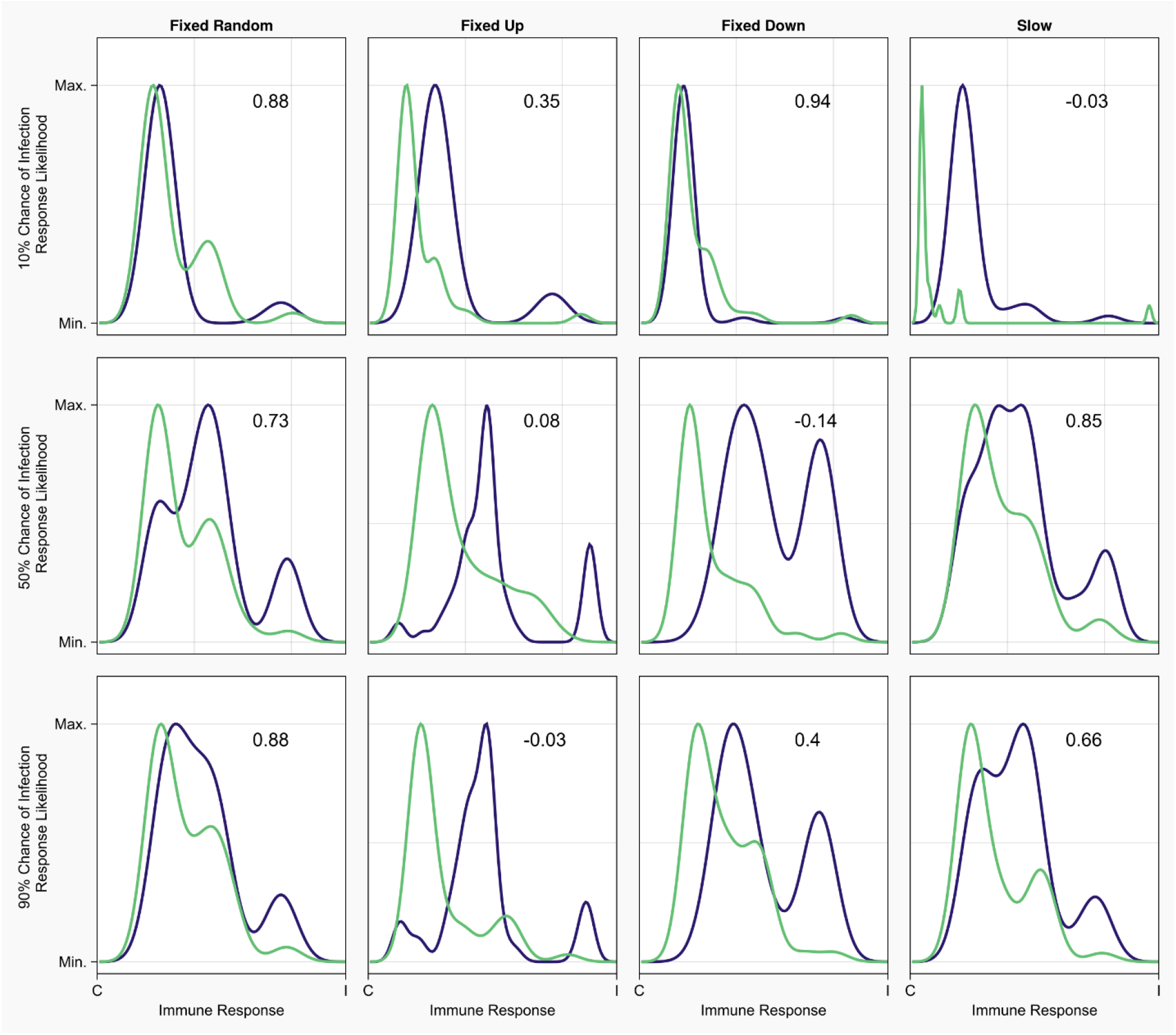
Immune response probability density function of pleiotropic winners (blue) and the last non-pleiotropic network (green) in the simulation. All plots on the same row have the same chance of infection (10%, 50%, or 90% descending), and plots in the same column have the same implementation of pleiotropy. 10 of 12 scenarios show pleiotropic winners being more inducible in their immune responses (as determined by right shifted density peaks) than their non-pleiotropic competitors. Each plot shows the results for competition after 250 generations of evolution.

### Pleiotropy alters immune signaling network evolution

Broadly, the immune responses generated by pleiotropic networks were identical to those generated by non-pleiotropic networks, not just in the distribution of their immune response on the constitutive-inducible spectrum but also in the peak amount of immune effector deployed during infection (Fig. 2A, B). When accounting for the differences in peak effector activity between constitutive and inducible immune responses, there was no difference in the magnitude of immune effector activity between pleiotropic and non-pleiotropic networks as a function of infection chance (Fig. 2B). In four scenarios, however (fixed upregulation when the chance of infection was 10% or 90%, fixed downregulation when the chance of infection was 50% or 90%), the Pearson correlation coefficient between the non-pleiotropic immune response probability density function and the pleiotropic immune response probability density function showed little to no correlation (*corr* ≤ ± .2), suggesting that these pleiotropic populations evolved significantly different immune responses than the non-pleiotropic populations.

When investigating immune effector level changes associated with signaling protein knockouts, in most conditions the loss of the pleiotropic protein did not result in significantly different effector activity relative to the removal of any other signaling protein (Fig. 3). Fixed downregulation is the only condition in which this was not true, indicating that theses hosts were reliant on the action of the pleiotropic signaling protein to produce their evolved immune response. On average, pleiotropic hosts were nearly as fit as non-pleiotropic hosts, with fixed downregulatory pleiotropy resulting in markedly higher absolute fitness than all other conditions when the chance of infection was 50% or 90% (Supplemental Fig. 3). Pleiotropic networks also had significantly different network sizes (Supplemental Fig. 4), connectivity (Supplemental Fig. 5), and distinct connections to effector proteins (Supplemental Fig. 6) depending on the pleiotropic element and the prevalence of parasites.

### Pleiotropy can imbue competitive benefits to organisms

We used two classes of competitive simulations to evaluate the fitness of the pleiotropic populations relative to the non-pleiotropic ones. Unevolved competition began immediately following host initiation and evolved competition began after host populations had evolved for 250 generations in isolation (i.e. in the absence of a different type of pleiotropic implementation). In unevolved competition, we observed that the pleiotropic upregulation element fixed in the population in most simulations when the chance of infection was 50% or 90%. Networks featuring pleiotropic downregulation fixed in the population when the chance of infection was 50%, while slowly evolving pleiotropic proteins had a high fixation rate at the intermediate infection risk. In evolved competition, networks featuring pleiotropic downregulation at high infection risk and those featuring fixed upregulation at intermediate infection risk fixed in most simulations. All results shown in Supplemental figure 1. Collectively, these results suggest that pleiotropic networks are capable of outcompete non-pleiotropic ones at intermediate and high infection risks over evolutionary time.

### Inducible immunity increases fitness relative to constitutive immunity, but is much rarer

The results that we obtained from this model generally favor the evolution of constitutive immunity as predominantly inducible immunity was rare in most scenarios, but it is unclear if this was due to high fitness imparted to hosts by constitutive immunity or the feasibility of producing an inducible immune response. The only populations in this work that consistently produced predominantly inducible immune responses where the pleiotropically downregulated hosts at higher infection risk levels (Fig. 2A). When comparing the absolute fitness of our populations we see that pleiotropic downregulation exceeded the fitness of non-pleiotropic hosts at higher infection risk levels, while all other network types were approximately equally fit (supplemental Fig. 3). This suggests that hosts expressing inducible immune responses are more fit than constitutive hosts. To investigate whether this absolute fitness advantage translated to a competitive advantage we looked at competitive simulations between pleiotropic and non-pleiotropic populations focusing on the most prevalent network from the winning population and the last network from the losing population (Fig. 4, Supplemental Fig. 2a-f). First comparing pleiotropic winners to non-pleiotropic losers we saw that in 10 of 12 scenarios inducible immunity was more common in the pleiotropic winners than in the non-pleiotropic losers (Fig. 4). Non-pleiotropic winners were similarly more inducible than pleiotropic losers (Supplemental 2 c). No pattern emerged when comparing pleiotropic winners and non-pleiotropic winners or pleiotropic losers and non-pleiotropic losers (Supplemental Fig. 2e, f). These results suggest that evolved inducible immune responses are more fit regardless of their interaction with pleiotropy, but that pleiotropy facilitates the evolution of inducible immunity which then bestows higher relative fitness on its host.

### Hosts initially express constitutive immunity and transition to inducible immunity

Examining the transition of response types over time could shed light on the network features that facilitate the evolution of predominantly induced immune responses. When looking at the immune responses and magnitude of response for each population in the first 50 generations of co-evolution we saw that in most scenarios populations rapidly converged on a single equilibrium over time, differing only in the magnitude of effector activated (sup. Fig. 8a-c). In the pleiotropically downregulated populations, however, we observed that populations branched toward multiple alternative states after converging on an early equilibrium. This difference was especially pronounced when the risk of infection was 90% (sup. Fig. 8c).

The degree of inducibility and magnitude of immune effector activated by hosts in the final generation did not seem to be restricted by the initial ancestor of a host. When looking at the lineages of hosts present in the last generation of a simulation, we found that hosts in inducible populations often shared a progenitor with hosts in constitutive or mixed-strategy populations (supplemental Fig. 9). While the complexity of our model prevents us from analytically solving for the existence of evolutionarily stable strategies, this behavior provides evidence for the presence of immune response strategies that are, at least temporarily, co-stable.

## Discussion

Our model of signaling network evolution allowed us to investigate the changes in evolutionary trajectories and endpoints that are associated with pleiotropic signaling proteins. Our findings suggest that pleiotropy in immune networks can be beneficial to organismal fitness, both by speeding the development of highly fit immune response dynamics and encouraging the exploration of phenotypic space by easing the transition from local fitness peaks to global ones. The effects of pleiotropy on organismal immune responses depend both on the specific action of the pleiotropic protein and on the prevalence of parasitic antagonists in the environment. These factors contribute to an evolutionary landscape where peak immune effector levels depend on parasite activity, and the mechanisms of achieving those peaks are heavily influenced by pleiotropic signaling proteins.

While the networks generated using this model cannot be explicitly related to any specific biological signaling pathway, there are important similarities between simulated host immune networks and those found in nature (supplemental fig. 7). In both cases, signaling networks make use of feedback loops between proteins and self-regulatory behavior is common (28,29). The evolved networks were sparsely connected, a feature often associated with biological signaling networks (30), with the mean connectedness of a population rarely exceeding 50% (supplemental Fig. 4). Some explicit parallels can be drawn to *Caernorhabditis elegans* immune defense, where distinct pathways with little crosstalk modulate immune activity (31), a feature observed in our populations as well (supplemental Fig. 7). We observed that our evolved immune networks mimic biological ones not only in structure but also in function; constitutively immune hosts have lower peak immune activity than hosts with inducible immune systems. This finding is consistent with expected stimulus responses in multilayered regulatory systems (32) and has been observed previously in explicit models of immune signaling networks (33). Our proposal that pleiotropic proteins can promote the evolution of complex traits, such as inducibility, is rooted in the understanding that extant biological processes are often co-opted when adaptive evolution gives rise to novel traits (21,23,34).

The hosts in our model tend to evolve constitutive immune responses, even when the chance of infection is low, a condition that has previously been hypothesized to favor inducible immunity (9,10,35). We believe this difference arises because we are not solely assessing the relative fitness of these immune strategies, but also their evolvability. The low abundance of parasites simply does not present enough of an evolutionary pressure for hosts to develop inducible immune networks. As parasite abundance increases, constitutive hosts carefully balance their effector activity against the rate of parasite replication to prevent growth, but rarely entirely remove parasites, in line with behavior predicted by Shudo and Iwasa (36). We see evidence of this balancing act in feedback loops between proteins that upregulate the effector and the effector itself, as well as effectors upregulating proteins that suppress their own activity (supplemental fig. 7). Constitutive immunity then represents a local fitness maximum that is easy to attain, especially for networks early in the evolutionary process (supplemental Fig. 8a-c).

Inducible immunity is generally a rare response in the context of our results, which is in line with previous work that modeled signaling networks explicitly (11) as well as those that developed analytical models of optimal trait usage (5,6,35). We might naively expect, then, that constitutive immunity would be more fit than inducible immunity. However, our results suggest that the opposite is true – populations that produced predominantly inducible responses were significantly more fit than their constitutive counterparts at higher infection risk levels (supplemental Fig. 3). When looking at population immune responses in the first 50 generations of a simulation, we see that hosts in the first generation predominantly mount constitutive responses across all conditions studied and that the first predominantly induced hosts do not appear until later generations (supplemental Fig. 8b, c). The scarcity of inducible immunity and the extended evolutionary time necessary for highly inducible immune responses to arise reinforces the notion that inducible immune responses are complex and evolutionarily challenging to deploy.

Besides being evolutionarily complex, highly inducible immune responses are more susceptible to disruption via parasites than constitutive immune responses, especially when parasite manipulation targets the pleiotropic signaling protein (Fig. 3), which may reinforce the prevalence of constitutive immunity (11). But, when a host with a predominantly inducible immune system does arise it is likely to rise to a high prevalence in the population. Collectively, our results paint a picture of inducible immunity being significantly more fit than constitutive immunity, especially as the magnitude of the induced response rises, with the scarcity of inducible behavior being explained by the evolutionary challenges to its development.

In conclusion, we have identified distinct changes in the trajectory of signaling network evolution associated with the inclusion of pleiotropic signaling proteins. Fixed random pleiotropy and evolutionary rate constraints on the pleiotropic protein did not result in significantly different evolved networks when compared to the non-pleiotropic control. Fixed upregulatory and fixed downregulatory pleiotropy altered initial and terminal network dynamics (supplemental Fig. 8a-c, Fig. 2), network size, connectivity, and the number of distinct paths from the detector to the effector (supplemental Fig. 4-6), all while maintaining mean population fitness that was approximately equal to or greater than non-pleiotropic hosts (supplemental Fig. 3a-c). With these findings we have provided some of the first evidence for the wide-ranging evolutionary effects of pleiotropic proteins on the signaling networks they are a part of, highlighting the importance of further empirical investigation into the benefits, tradeoffs, and evolutionary consequences of pleiotropy in the immune system and across the genome. This work advocates for using a broad perspective when studying known pleiotropic proteins and genes, as their full evolutionary effects may only be observed at the scale of signaling networks.

## Materials and Methods

The model of signaling network evolution that we based this work on was originally designed model the evolution of generic signaling networks (37), and was subsequently modified to study immune signaling network robustness (38) and the evolvability of immune responses during coevolution (11). Here we have revised the model to include pleiotropic signaling proteins. We designed the following implementations of pleiotropy: 1) random connections between the pleiotropic protein and other proteins in the network that are fixed across evolutionary time; 2&3) a single connection from the pleiotropic protein that up-or downregulates the effector as a side effect of its function in development; 4) pleiotropic proteins that may evolve but undergo reduced mutation rates relative to other network components, capturing reduced evolutionary rates associated with purifying selection. Figure 1 provides a diagrammatic representation of these restrictions compared to the non-pleiotropic case.

We used two broad classes of simulation to study the effects of pleiotropy on immune evolution: co-evolution and competition. In co-evolution simulations, a population of hosts evolved for 500 generations with a population of parasites. The host immune networks in these simulations were either non-pleiotropic or all hosts in the population had the same pleiotropic constraint as defined above. Each simulation had 500 hosts, with each host initially defined by a randomly generated immune network. Parasite population size was determined by the chance of infection in each simulation, and each parasite possessed a single connection to a signaling protein that could disrupt host immune signaling. We used these simulations to study immune networks and their dynamics when evolving under pleiotropic constraint.

Competitive simulations were broken into two phases: independent evolution and competition. During independent evolution a non-pleiotropic population (n = 250) and a pleiotropic population (n = 250) were allowed to co-evolve with parasite populations without interacting with the other population of hosts. After 250 generations the simulation entered competition, combining the host populations along with their parasites. The competition ended when one population died out entirely or 1000 generations had passed with no winner (draw). These simulations allowed us to evaluate the viability of pleiotropy in a population that is not uniformly facing the same potential fitness deficits. In all cases, for each implementation of pleiotropy and chance of infection, we conducted 100 simulations.

Each host network initially contains a single detector, three signaling proteins, and a single effector. Over the course of a simulation, mutations during reproduction duplicated or deleted signaling proteins and deleted, added, or altered regulatory interactions between proteins in the network. Hosts remained restricted to a single detector and a single effector, and at no point were detectors and effectors allowed a direct connection. Host fitness was determined as a function of immune effector activity pre- and post-infection, cumulative parasite load, and network size (see equation 3). Parasite fitness was strictly based on cumulative parasite load during infection, a proxy for transmission potential.

### Simulation Framework

Evolutionary simulations were carried out in the Julia programming language (v 1.6) and can be broken into the following discrete events.

#### 1. Initialization

a population of hosts is generated at random. All hosts start with a detector, three signaling proteins, and an effector. Each possible connection in the network is given a 50% chance of occurring, and for each connection that does occur a random number is selected on the interval [−1,1] to decide the regulatory behavior of that connection. The only constraint on initial network structure is that the detector protein cannot directly connect to the effector protein.

A population of parasites is also generated to infect 10%, 50%, or 90% of hosts, depending on the chance of infection for the simulation. Parasites are designated by a target (restricted to signaling proteins) and regulatory behavior on that target (restricted to the interval [−1,1]).

#### 2. Healthy Equilibrium

Each protein P in an immune network has total concentration equal to 1, including an active portion *P*_i_^*^ and inactive portion *P*_*i*_, ([P_i_^*^] + [*P*_*i*_] = 1). During infection, changes in parasite abundance are calculated as though it was another protein in the network. The change in 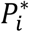. R is determined by the regulatory action on *P*_*i*_ defined:

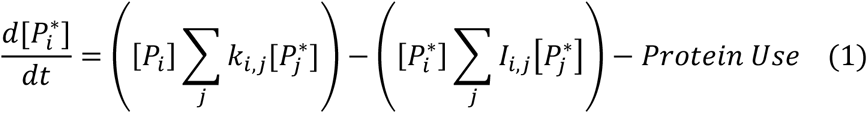

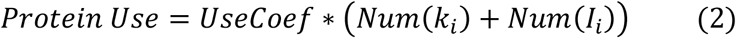

Where *k*_*i,j*_ are the upregulatory coefficients from protein *P*_*j*_ to protein *P*_*i*_R, *I*_*i,j*_ are the downregulatory coefficients from protein *P*_*jj*_ to protein *P*_*i*_. The term *Protein Use* describes the inactivation of some 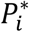 owing to the action of *P*_*ii*_ on other proteins in the network. Starting from initial concentrations of *P*^*^ = .5 for all proteins in the network, *P*^*^ for each protein at time step *t+1* is calculated numerically using equation (1) until the effector protein reaches equilibrium.

#### 3. Infection

based on the chance of infection, l0%, 50%, or 90% of hosts are randomly selected for infection by parasites. Parasites are treated as an additional protein in the host immune network with an upregulatory connection of 1 to the host detector, a self-targeted upregulatory connection of .8 to represent replication, and an up-or downregulatory connection to a signaling protein targeted by the parasite. The immune effector of the host network gains a downregulatory connection of −1 directed toward the parasite; see figure 1 for an example infection.

#### 4. Infected Dynamics

Equation (1) is used to calculate changes in protein and parasite levels until one of the following occurs:

A. Twenty time-steps have passed, which we consider the end of the hosts reproductive lifespan
B. Parasite concentration drops below 1e-2, where the parasite is considered cleared, or
C. Parasite concentration exceeds .9, where the host succumbs to the infection.

#### 5. Fitness Calculation

Host fitness is calculated using the following equation:

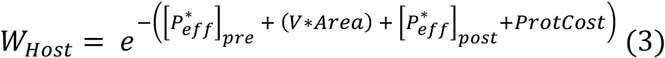

With

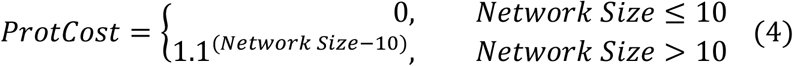

The variable *V* is parasite virulence, *Area* is the area under the parasite infection curve normalized to 1, and *Network Size* is the number of proteins in the network. The network size cost used the size of the toll pathway as a neutral benchmark (39). The increase in cost with increasing size reflects the fitness deficits associated with maintaining and producing large amounts of proteins (24). Parasite fitness was taken strictly from the normalized area of the parasite infection, which is representative of the total number of parasites produced over the course of the infection.

#### 6. Host and Parasite Death

in each generation up to 30% of each population dies, with hosts that succumbed to infection and parasites that were cleared by their hosts at the top of the priority list for death. In the case where greater than 30% of either population was removed in this manner, individuals selected for death were randomly added back to the healthy population 70% of individuals survived infection.

When less than 30% of the population died in the infection phase, individuals were selected to die in a fitness weighted manner. At random an individual was selected from the population and its chance of dying was inversely proportional to its relative fitness against the population. Ex: a host that was in the 75^th^ percentile of fitness has a 25% chance of dying. Individuals were selected until 30% of the population died or until each had been tested in this manner.

#### 7. Host Reproduction

survivors are picked at random to reproduce, completely replenishing the host population (keeping population size constant across generations). Reproduction either results in the creation of a direct copy of the parent or, rarely, a mutated copy (host mutation rate: 5e-3). For hosts, a mutation could be one of the following modifications to the parent network:

1. Add a protein-protein interaction between two randomly selected proteins with a random regulatory behavior (relative probability = .25)
2. Delete a protein-protein interaction (relative probability = .25)
3. Alter regulatory coefficient (relative probability = .3)
4. Delete a protein (relative probability = .1)
5. Duplicate a protein (relative probability = .1) ^*^

^*^Duplication is the only mutation that can affect the pleiotropic signaling protein. The duplicated pleiotropic protein is treated as a non-pleiotropic signaling protein

#### 8. Parasite Reproduction

surviving parasites are picked to reproduce in a purely fitness-based manner, with highly fit parasites producing more offspring than their lower fitness peers. This corresponds to higher cumulative parasite load leading to more offspring in the following generation. Parasites with a cumulative load between 0 and .33 produced one offspring, those with a cumulative load between .34 and .66 produced two offspring, and finally a cumulative load between .67 and 1 lead to 3 offspring. Parasites reproduced until the population is completely replenished. Parasites reproduced by way of direct copy of the parent, or rarely a mutated copy was produced (mutation rate of 1e-2 for parasites). Parasite mutations are defined as follows:

1. randomly change the target signaling protein (relative probability = .5)
2. change the regulatory behavior on the targeted signaling protein to a new value in [−1,1] (relative probability = .5)

After step 8, the simulation repeats starting at step 2 using the hosts and parasites populations produced in step 7 & 8. Each of these cycles from step 2-8 is termed a generation and each simulation runs for 500 generations. For each combination of infection rate and pleiotropic constraint we ran 100 simulations. This number of simulations was chosen to balance computational time against reproducibility.

### Competitive Simulations

We devised competitive simulations to determine the relative fitness differences between pleiotropic and non-pleiotropic hosts in this model. We conducted 100 competitive simulations for each implementation of pleiotropy and chance of infection pairing, and these simulations had the following changes from the non-competitive case described above:

1. 250 pleiotropic hosts and 250 non-pleiotropic hosts either immediately entered competition (unevolved competition) or were allowed 250 generations to evolve independently (evolved competition), at which point their populations were merged and they entered competition.
2. Competitive simulations proceeded until one of the host populations was extinct, resulting in a victory for the extant population, or until 1000 generations had passed with no winner, resulting in a draw.

### Data Analysis

#### Immune Response Probability Density

To study the immune responses mounted by host populations, we first calculated the percentage of each host’s maximal immune response that was activated by infection. For each host this generates a value describing the degree of inducibility of their immune response, with 0% indicating no induced response to parasites, and 100% indicating an entirely induced response. We then approximated the probability density function for this data using kernel density estimation. This immune response probability density conveys the likelihood that a host in a population would have a specific percentage of their immune response induced by parasites and these values were normalized to one to ease comparisons between populations. All infected hosts in the final generation of each simulation were used in the calculation of immune response probability density functions.

#### Correlation

We calculated the Pearson correlation coefficient to aid in rigorous comparisons between pleiotropic and non-pleiotropic host immune response densities. For each combination of pleiotropic implementation and chance of infection, we calculated the Pearson correlation coefficient between pleiotropic immune response density and non-pleiotropic immune response density at the same chance of infection. Incidents of poor correlation (*corr* ≤ |.2|) suggest a significant difference between the immune responses mounted by the pleiotropic and non-pleiotropic populations.

#### Immune effector activity vs immune response type

To visualize the relationship between immune response type and peak immune effector activity we calculated a two-dimensional probability density function, where the x axis was the proportion of the response induced by parasite and the y axis was the maximum amount of immune effector activated. These probability density functions were calculated for each implementation of pleiotropy and chance of infection pairing using kernel density estimation. All hosts infected in the final generation of each simulation were used to generate these plots. The two-dimensional probability density functions for each pairing of pleiotropic implementation and chance of infection were then added together to produce Figure 2B.

#### Determining the effect of signaling protein knockouts

To determine if pleiotropic hosts were more susceptible to manipulation than non-pleiotropic hosts, we calculated the mean absolute difference in active effector levels between intact hosts and hosts with a single signaling protein knockout (the protein was removed from the network). Hosts were infected with a non-disrupting parasite for twenty time-steps and the effector levels for the intact network and the knockout for each signaling protein were measured. The mean of the absolute difference in effector levels at each time step between the intact and knockout networks was calculated and is used as a metric of the networks reliance on a specific signaling protein to produce the evolved response. Significant differences between the mean absolute difference in effector level following knockout of the pleiotropic signaling protein was compared that of non-pleiotropic knockout using two-tailed homoscedastic t-tests with Bonferroni correction.

#### Network Features

Network connectivity was calculated by dividing the number of protein-protein interactions in a network and dividing that number by the number of possible connections that network could possess. To determine if pleiotropy altered the number of proteins necessary to mount an immune response, we measured network size by counting the number of proteins present in the most common immune network from the end of each simulation for a given implementation of pleiotropy and chance of infection. Finally, as a measure of robustness, we calculated the number of distinct paths from the detector to the effector. A distinct path was one that did not share a signaling protein with any other path connecting the detector and effector, and previous empirical work has proposed this to be a mechanism promoting robustness to parasitic interference in immune network signaling (31).

## Code Availability

All code used to generate the results of this model are available on Github (https://github.com/Reese-Martin/Immune_Evolution_and_Pleiotropy) and will be packaged and archived with a permanent DOI on Zenodo upon acceptance of the manuscript.

## Supporting Information Captions

**Supplemental Table 1:** Names, values, and description for variables and parameters used in the simulation

**Supplemental Figure 1:** Results of competition simulations.

**Supplemental Figure 2:** Winners and losers of competitive simulations after 250 generations of adaptation

**Supplemental Figure 3:** plots of average host fitness through 500 generations.

**Supplemental Figure 4:** Size (number of proteins) of the most common network from each run of a scenario.

**Supplemental Figure 5:** Percentage of total potential connections deployed by the most common network at the end of each simulation.

**Supplemental Figure 6:** Number of distinct paths from the detector to the effector in the most common network following a simulation.

**Supplemental Figure 7:** The average host network generated in each pleiotropic constraint and infection level pairing.

**Supplemental Figure 8:** Magnitude of immune response by the proportion of the response that is induced in the initial 50 generations

**Supplemental Figure 9:** The proportion of runs that ended in a lineage bifurcation (multiple discrete immune strategies descended from a single lineage)

