## Supplementary Information for "Pleiotropy promotes the evolution of inducible immune responses in a model of host-pathogen coevolution"

**Supplemental Table 1:** Names, values, and description for variables and parameters used in the simulation

| Variable | Value | Description |
|---|---|---|
| Parasite damage ($v$) | 2 | The amount of damage incurred by hosts due to parasite infection, akin to virulence. Chosen so that excess immune activity and parasite burden had the same degree of effect on fitness |
| DeathCoef | .3 | The percentage of each population that died in a single generation |
| DeathThreshold | .9 | The limit of parasite burden that hosts can tolerate before being unable to breed |
| UseCoef | .01 | The amount of protein $P_i$ that is deactivated in a single time step due to acting on protein $P_j$ |
| Host Mutation Rate | 5e-3 | Selected to be balance scarcity of mutations found in nature against needing to run simulations for more generations. Picked in coordination with parasite mutation rate so that parasites mutate at 2x the host rate |
| Parasite Mutation Rate | 1e-2 | Selected to be balance scarcity of mutations found in nature against needing to run simulations for more generations. Picked in coordination with parasite mutation rate so that parasites mutate at 2x the host rate |

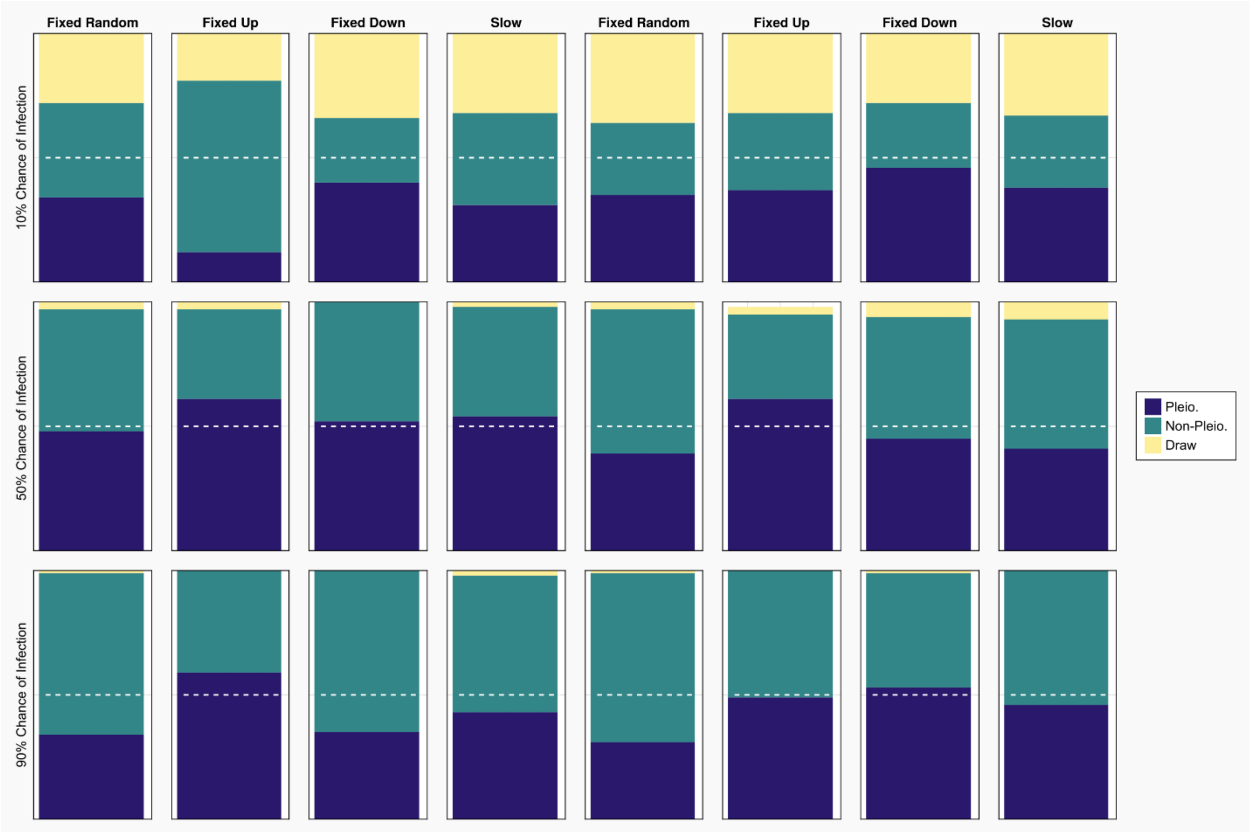


**Supplemental Figure 1:** Results of competition simulations. Rows correspond to infection percentages and columns correspond to the pleiotropy type for a set of competitions. Unevolved competitions are those that had non-pleiotropic and pleiotropic organism enter competition immediately. Evolved are those that took place after 250 generations of adaptation in isolated populations.

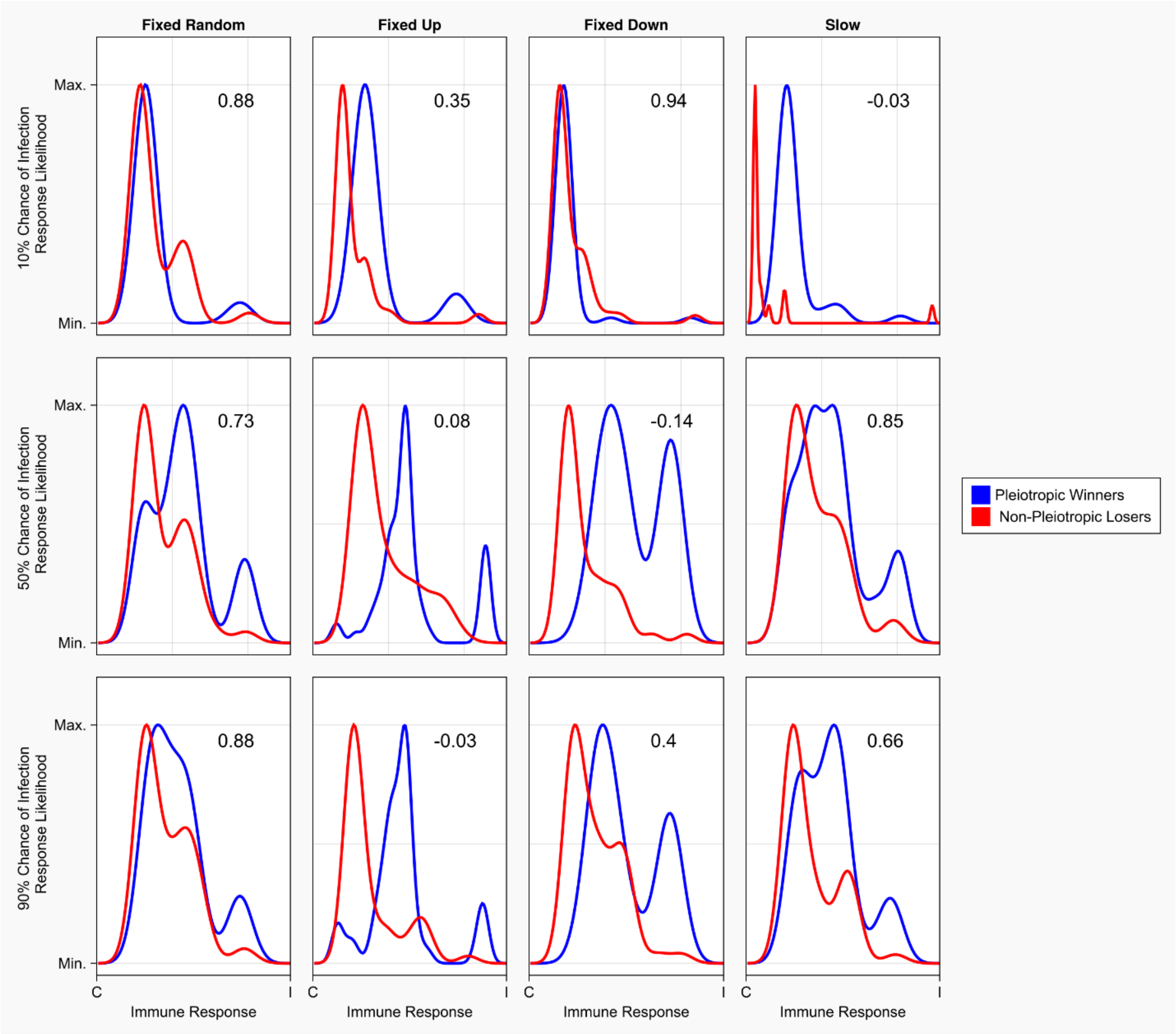


**Supplemental Figure 2a:** Winners and losers of competitive simulations after 250 generations of adaptation: Pleiotropic winners (blue) vs Non-pleiotropic losers (red).

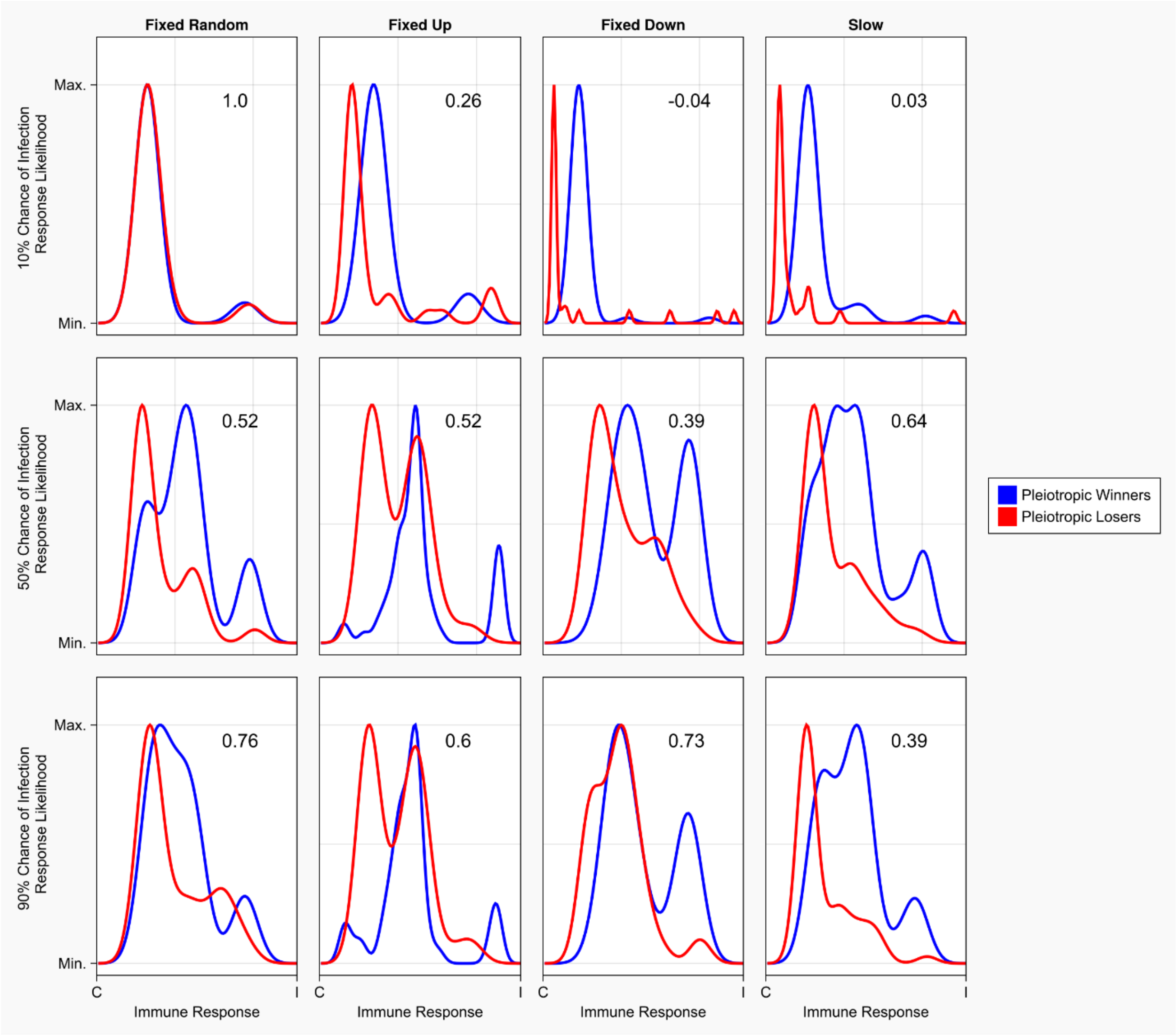


**Supplemental Figure 2b:** Winners and losers of competitive simulations after 250 generations of adaptation**:** Pleiotropic winners (blue) vs Pleiotropic losers (red).

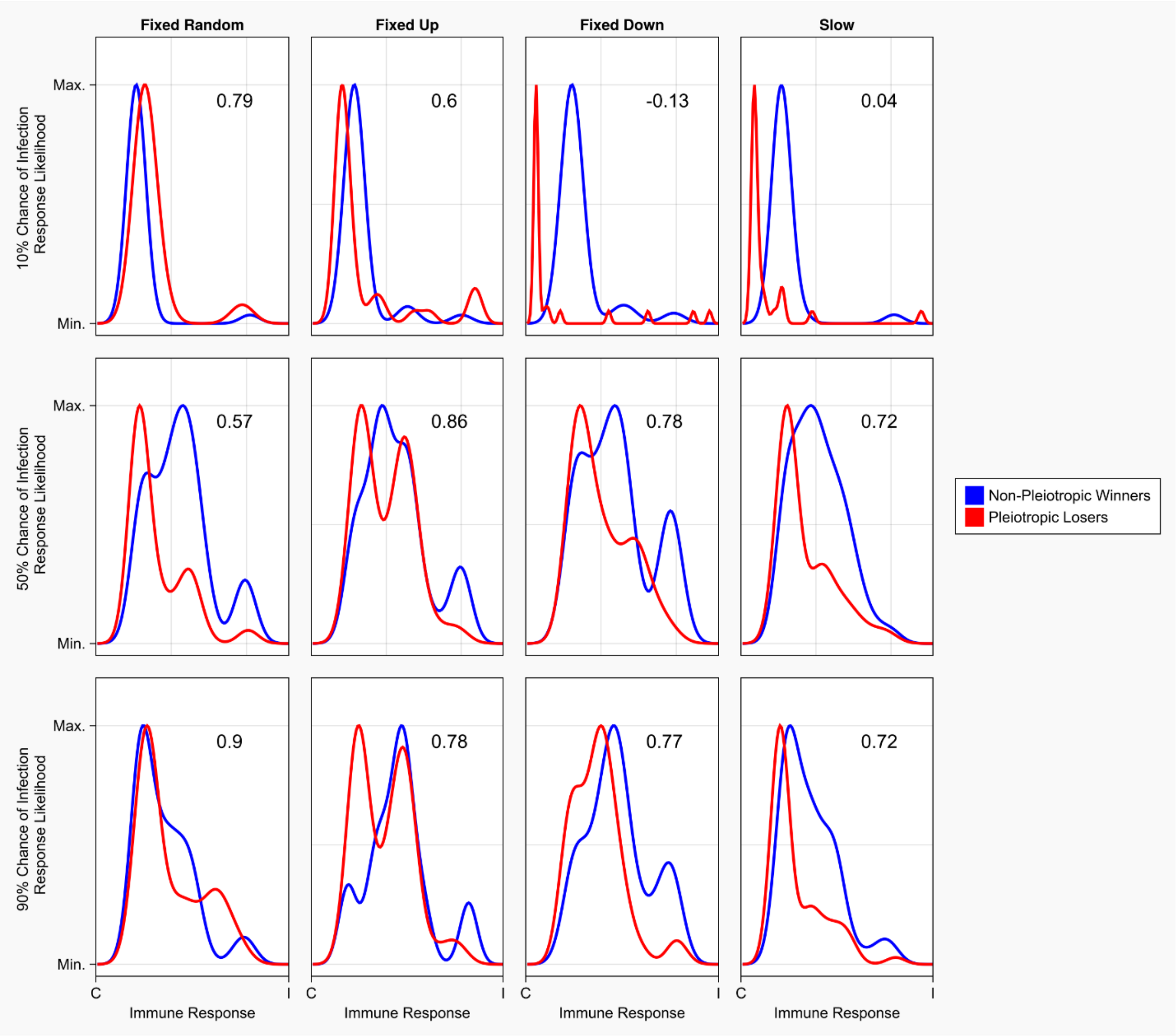


**Supplemental Figure 2c:** Winners and losers of competitive simulations after 250 generations of adaptation**:** Non-pleiotropic winners (blue) vs Pleiotropic losers (red).

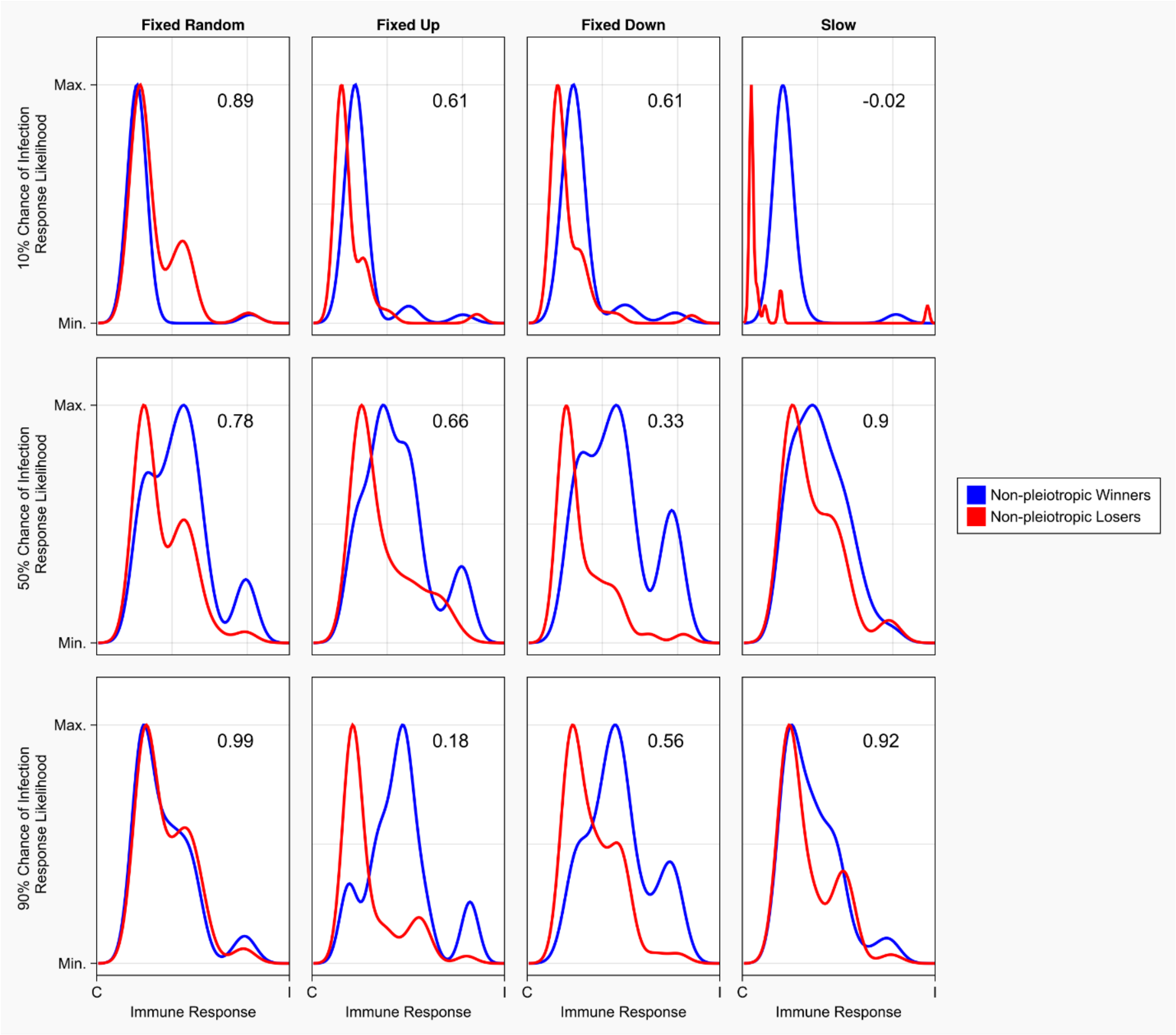


**Supplemental Figure 2d:** Winners and losers of competitive simulations after 250 generations of adaptation**:** Non-pleiotropic winners (blue) vs Non-pleiotropic losers (red).

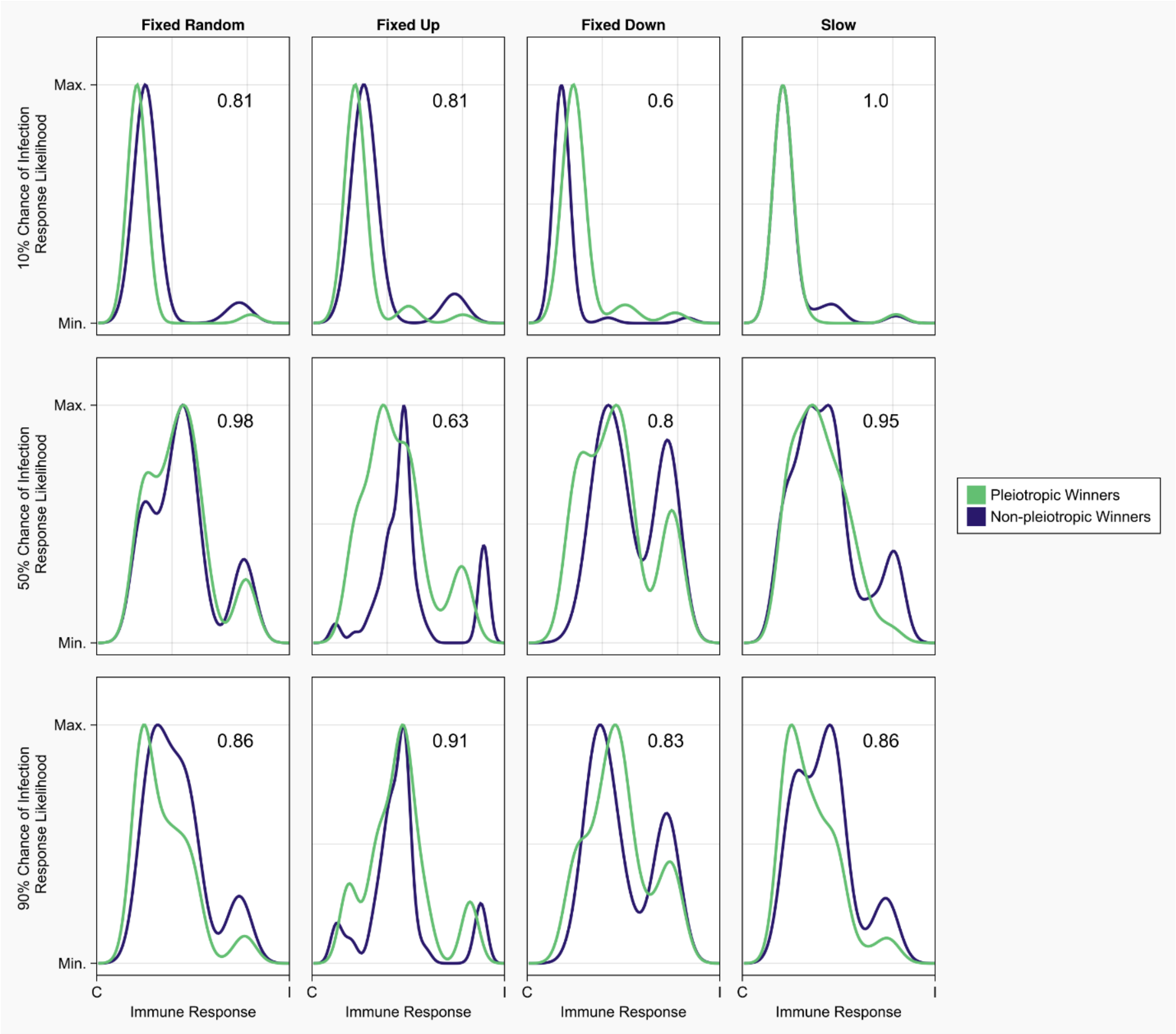


**Supplemental Figure 2e:** Winners and losers of competitive simulations after 250 generations of adaptation: Pleiotropic winners (blue) vs Non-pleiotropic winners (green).

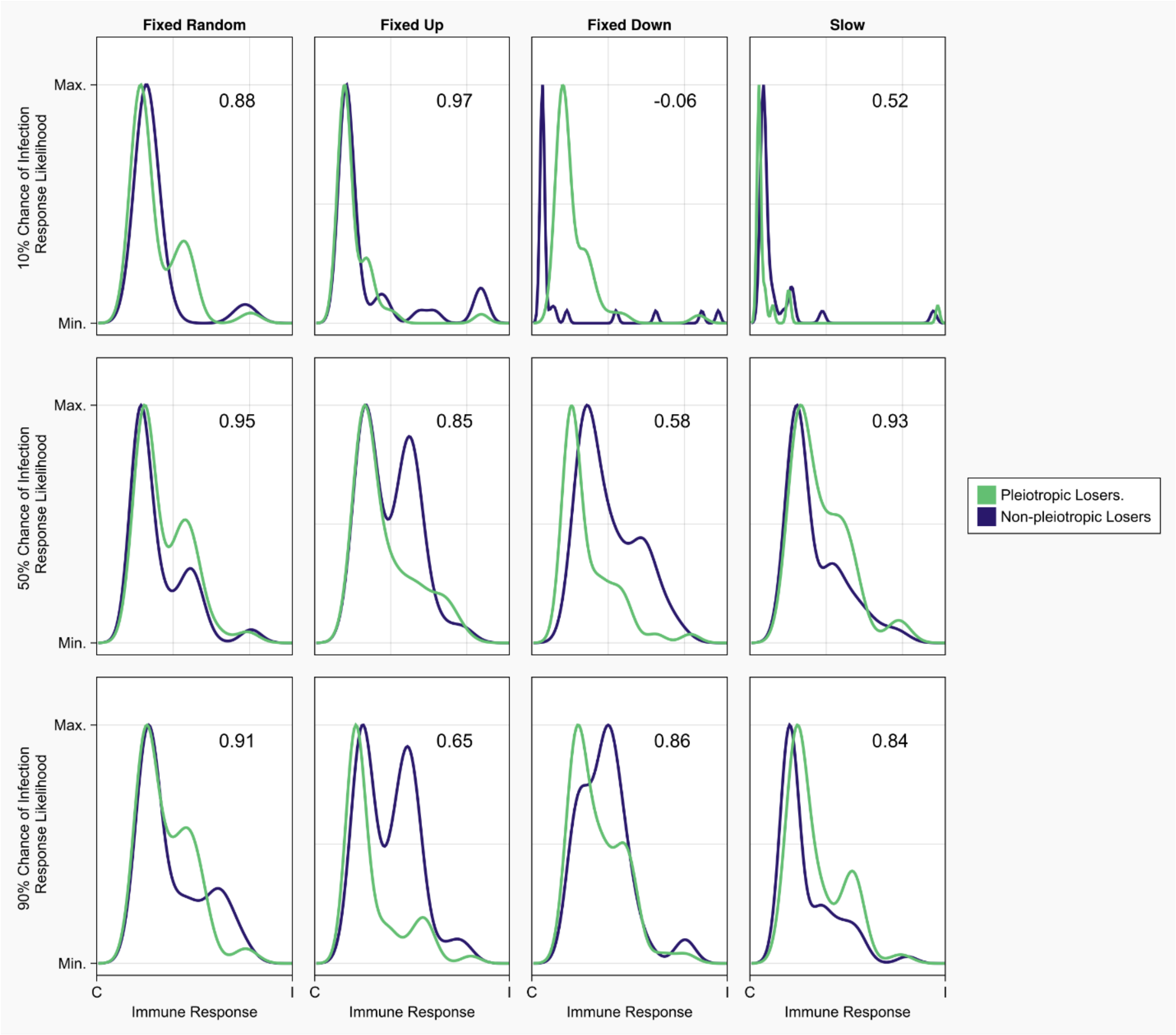


**Supplemental Figure 2f:** Winners and losers of competitive simulations after 250 generations of adaptation**:** Pleiotropic losers (blue) vs Non-Pleiotropic losers (green).

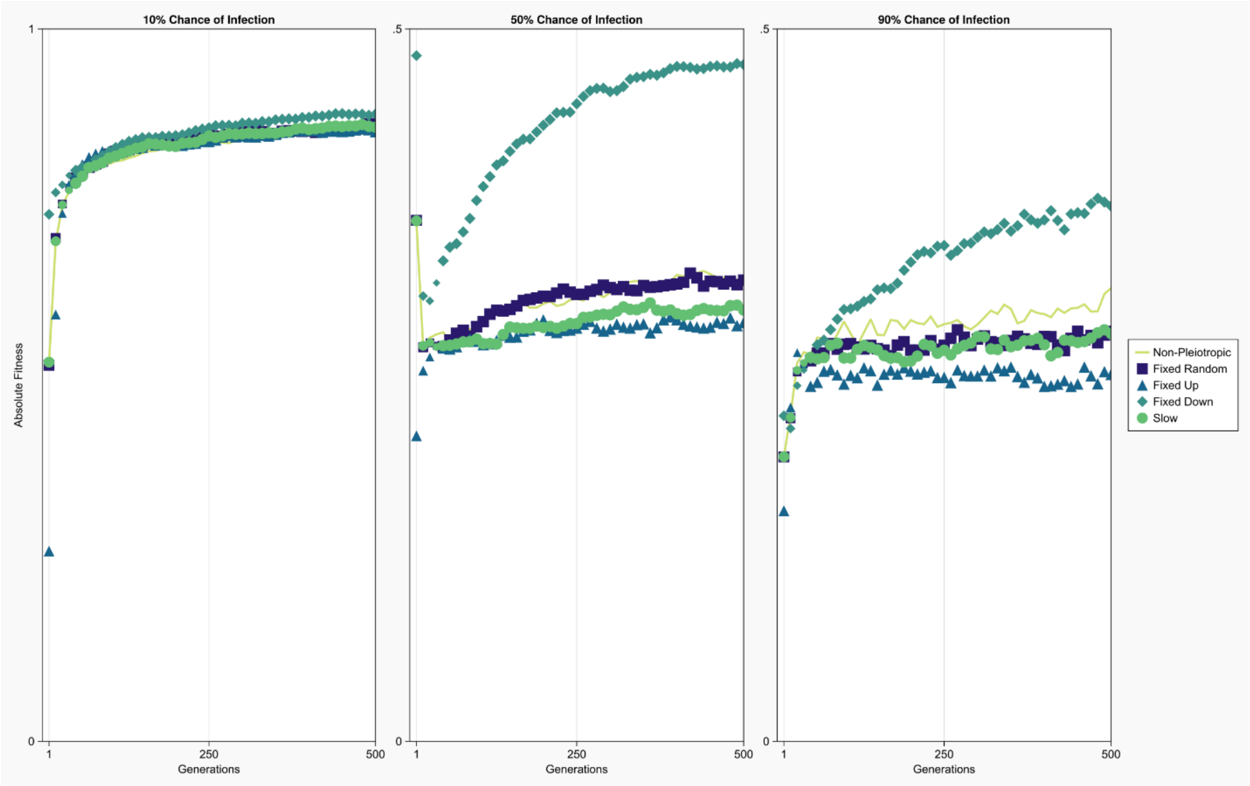


**Supplemental Figure 3:** plots of average host fitness through 500 generations when the chance of infection was 10%( left), 50% (middle), or 90% (right). Each panel shows host or parasite fitness from unconstrained (solid line), Fixed Random (squares), Fixed Up (triangles), Fixed Down (diamonds), and 100x slower evolution (circles) simulations. Average host fitness was calculated using hosts that were and were not infected for each generation. Note that the y axis changes scale in the second and third panel to because overall host fitness decreased as the chance of infection increased.

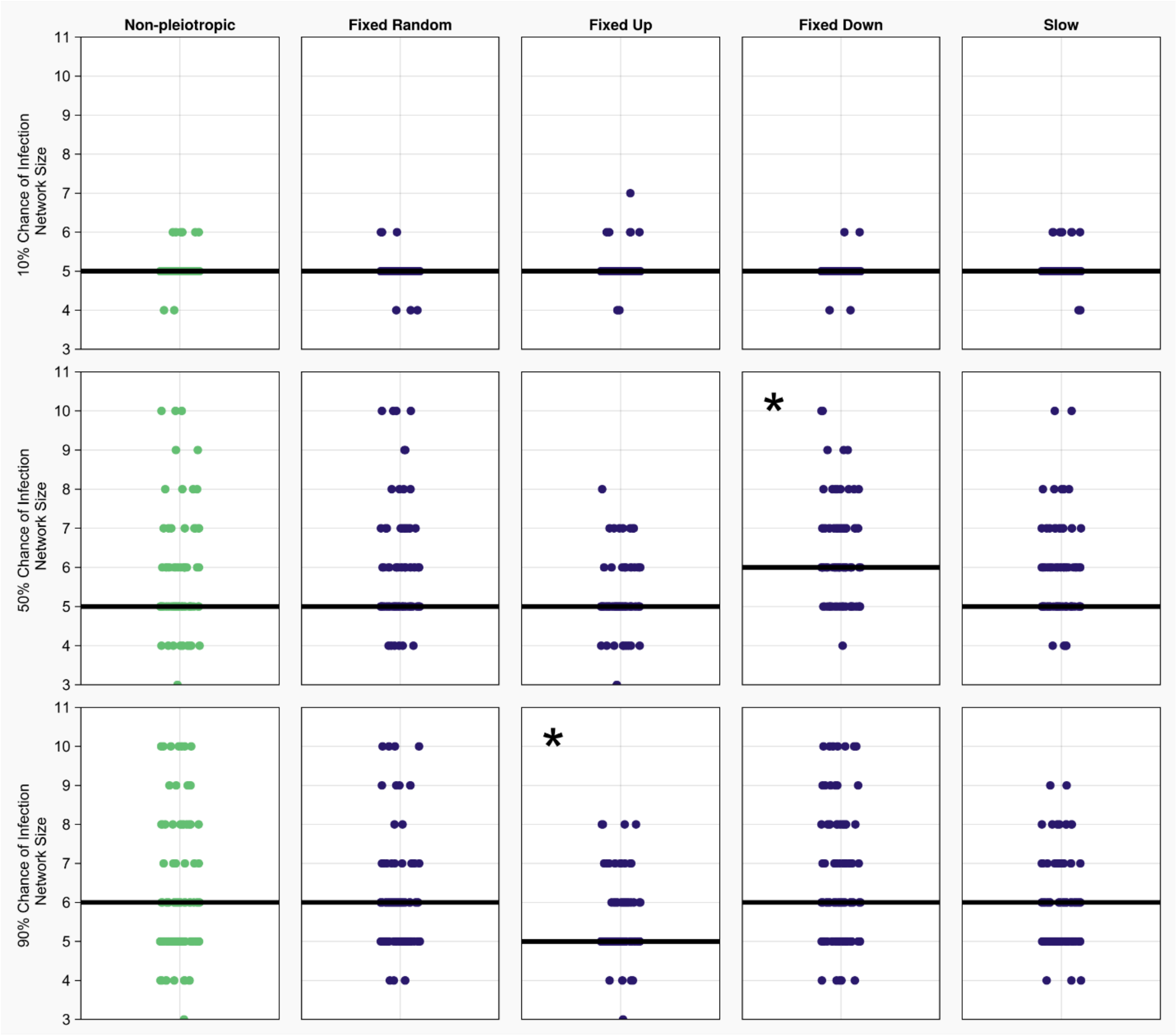


**Supplemental Figure 4:** Size (number of proteins) of the most common network from each run of a scenario with median lines presented in black. All networks start with 5 proteins, and at the 10% infection rate, no pleiotropic conditions diverge significantly from this point. At the 50% infection level, only the fixed downregulation pleiotropic condition has an increased median network size. Conversely, at the 90% infection level, only fixed upregulation did not increase in median network size. Asterisks indicate a significant difference from the non-pleiotropic scenario in each row.

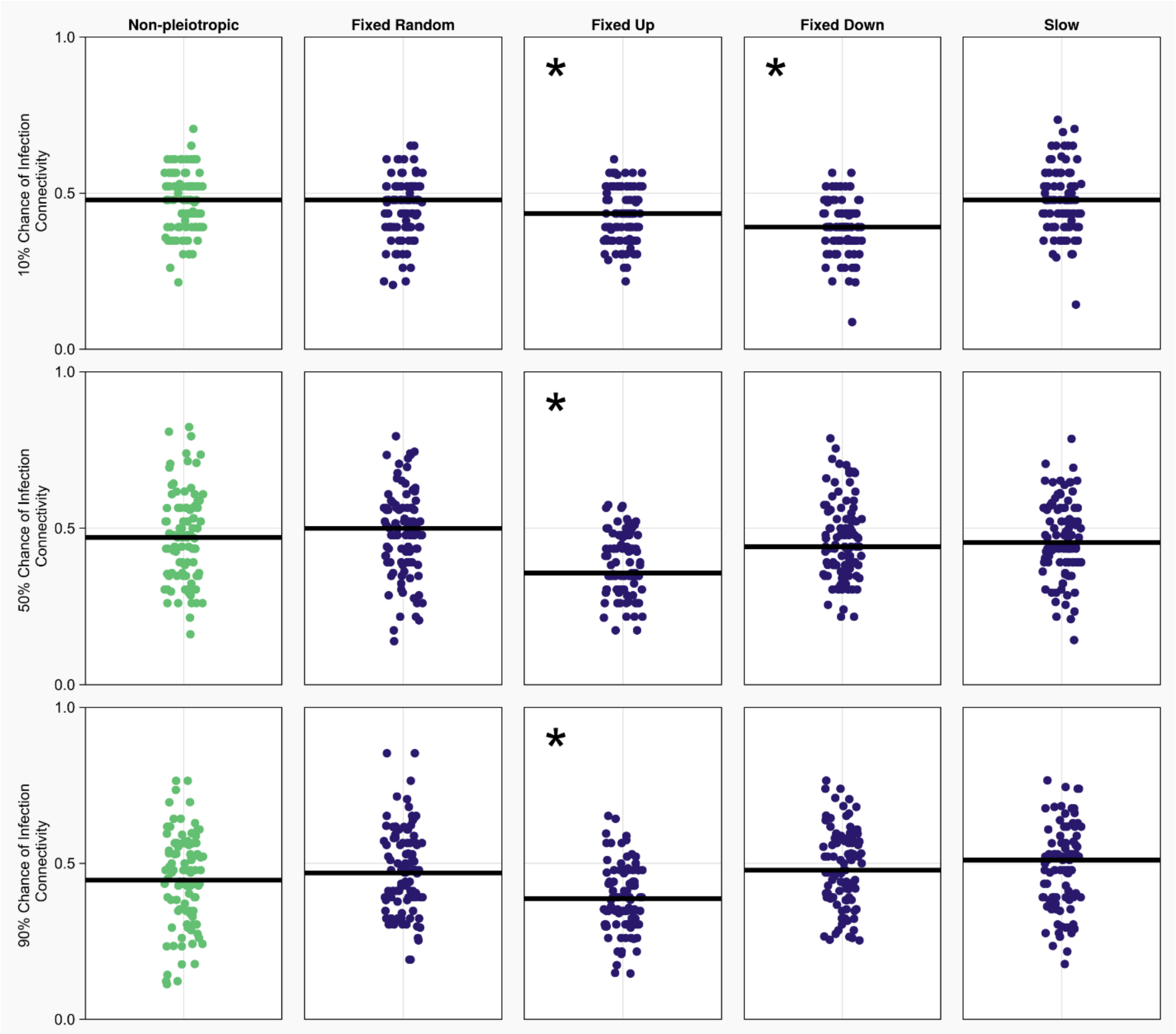


**Supplemental Figure 5:** Percentage of total potential connections deployed by the most common network at the end of each simulation. Each row is an infection risk (10, 50, or 90%) and each column is a type of pleiotropy. On average, half of all connections are used initially. Asterisks indicate a significant difference from the non-pleiotropic scenario in each row.

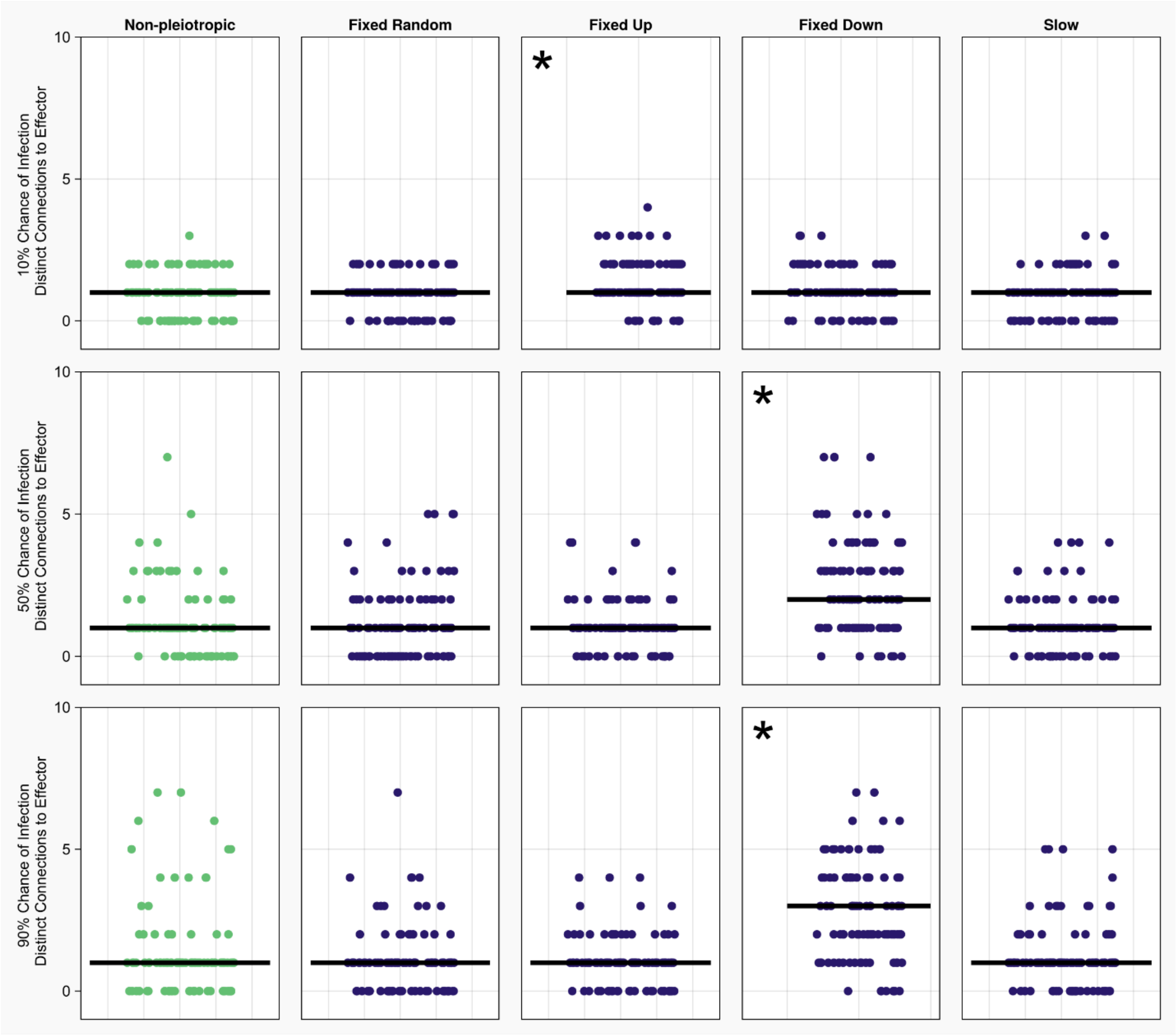


**Supplemental Figure 6:** Number of distinct paths from the detector to the effector in the most common network following a simulation. The distinct paths through a network are the set of paths that share no signaling proteins with the other paths in the set. The fixed downregulation conditions deploy a higher number of distinct paths. Importantly, distinct paths connect the detector to the effector in a manner that is partially insulated from other paths through the network, increasing robustness. Asterisks indicate a significant difference from the non-pleiotropic scenario in each row.

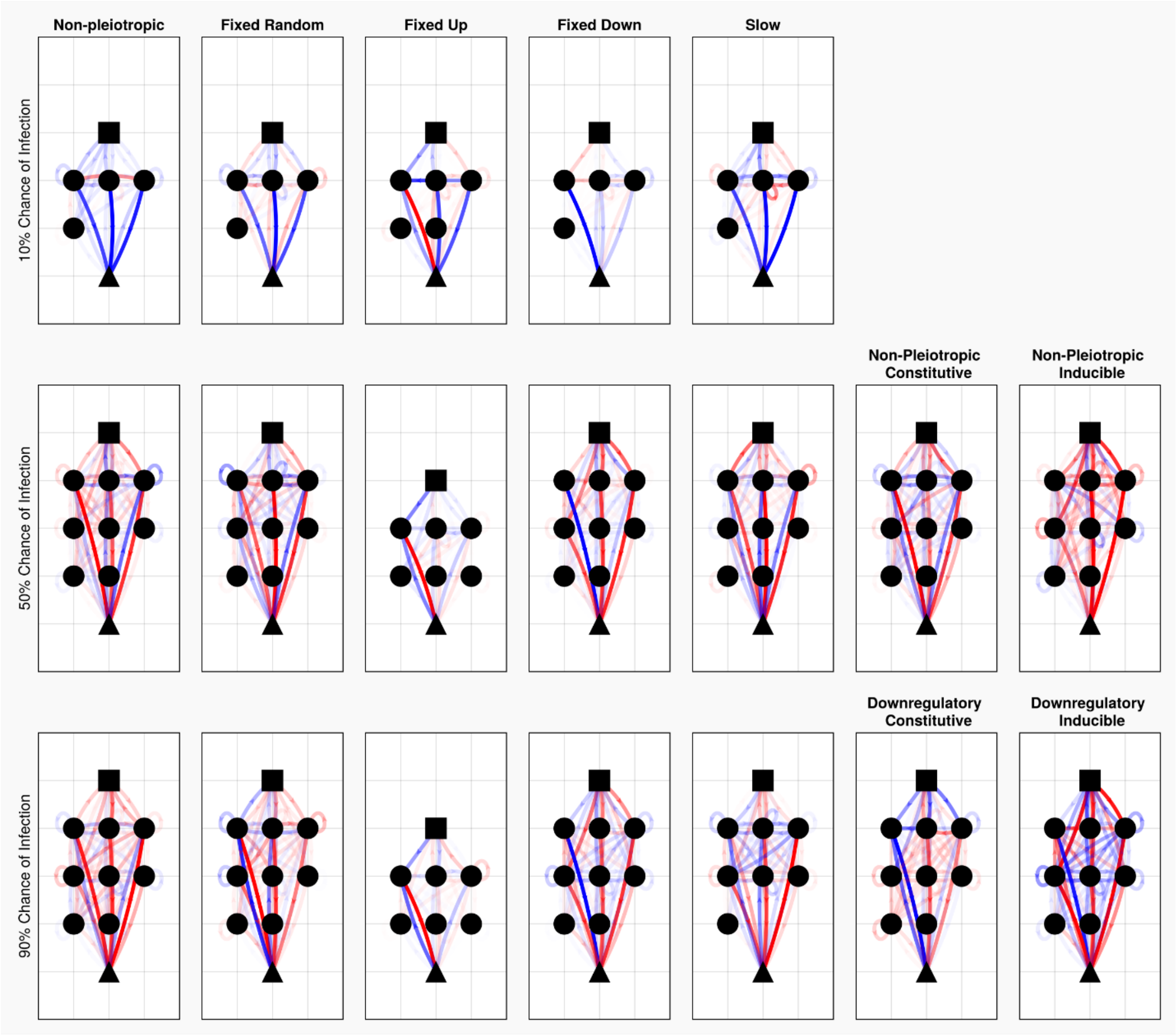


**Supplemental Figure 7:** The average host network generated in each pleiotropic constraint and infection level pairing. These average networks were generated using the most common networks from the end of each simulation at a given pairing. Saturation for the connections between proteins is scaled based on the most common connection across all networks at the given constraint and infection level. Arrows denote the direction of the connection, blue connections are down regulatory, red are upregulatory.

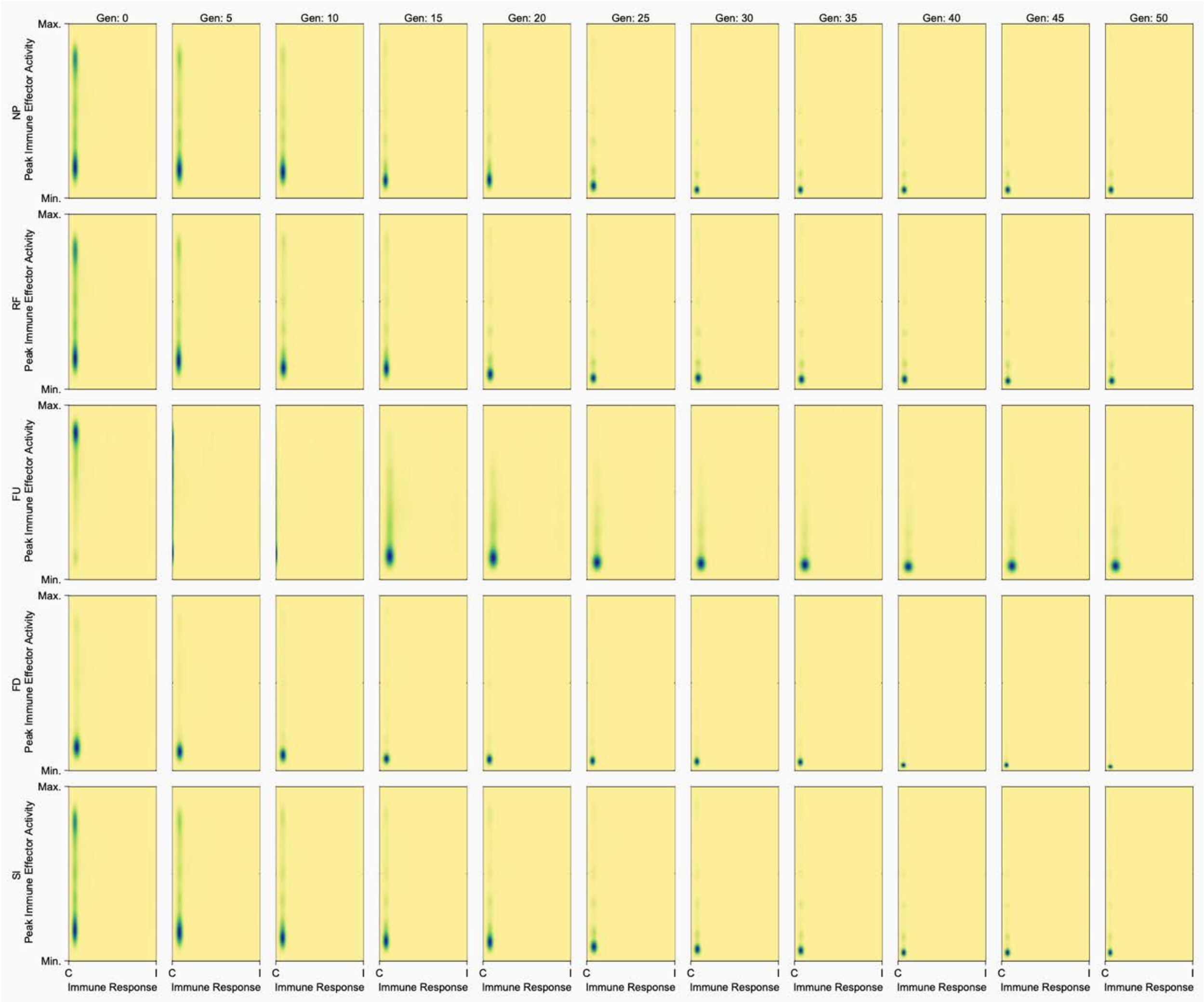


**Supplemental Figure 8a:** Magnitude of immune response by the proportion of response that is induced in the initial 50 generations of an evolutionary simulation where the chance of infection was 10%. Darker colors indicate more common combinations of magnitude of immune responses and proportion of response induced by parasites.

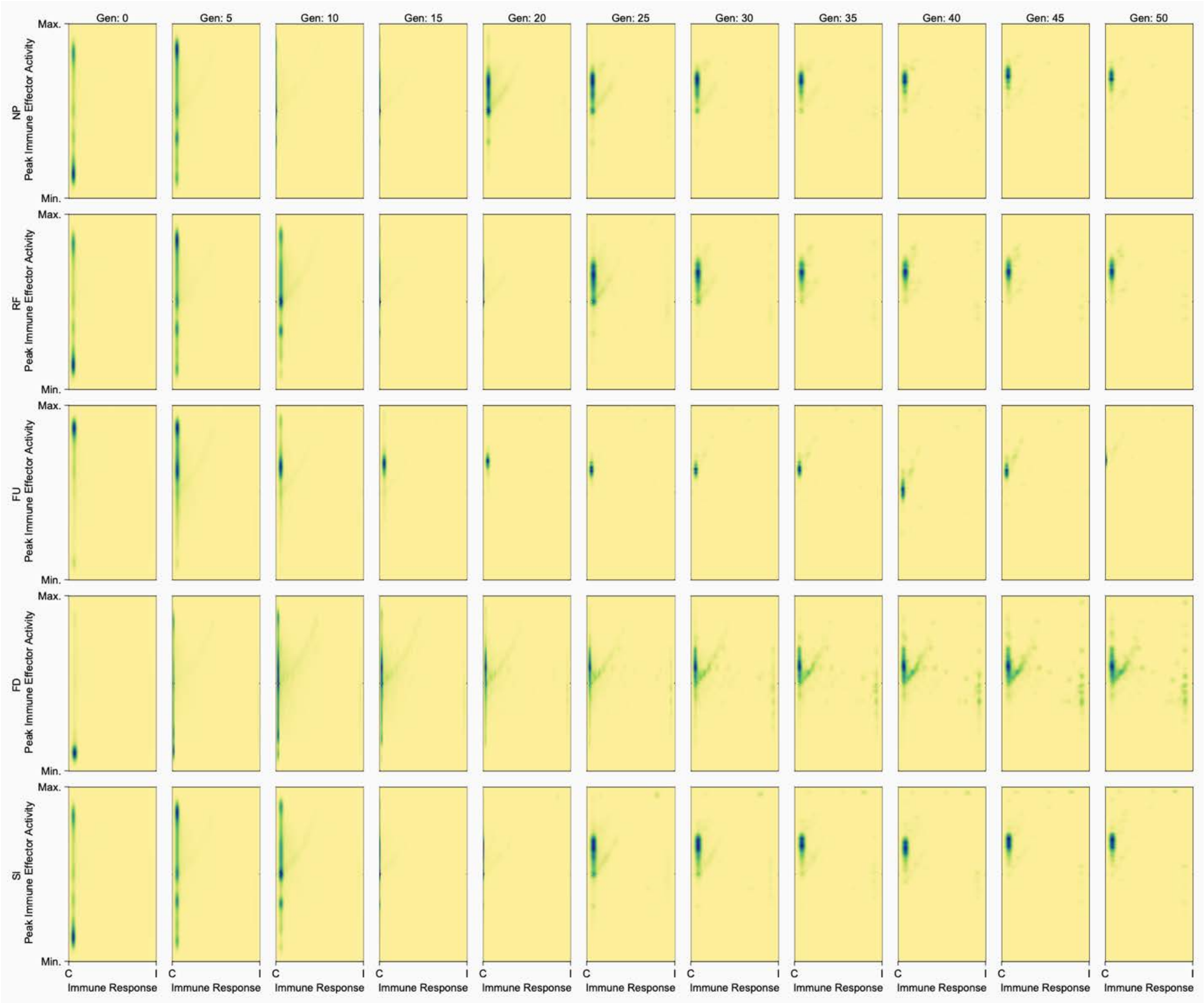


**Supplemental Figure 8b:** Magnitude of immune response by the proportion of response that is induced in the initial 50 generations of an evolutionary simulation where the chance of infection was 50%. Darker colors indicate more common combinations of magnitude of immune responses and proportion of response induced by parasites.

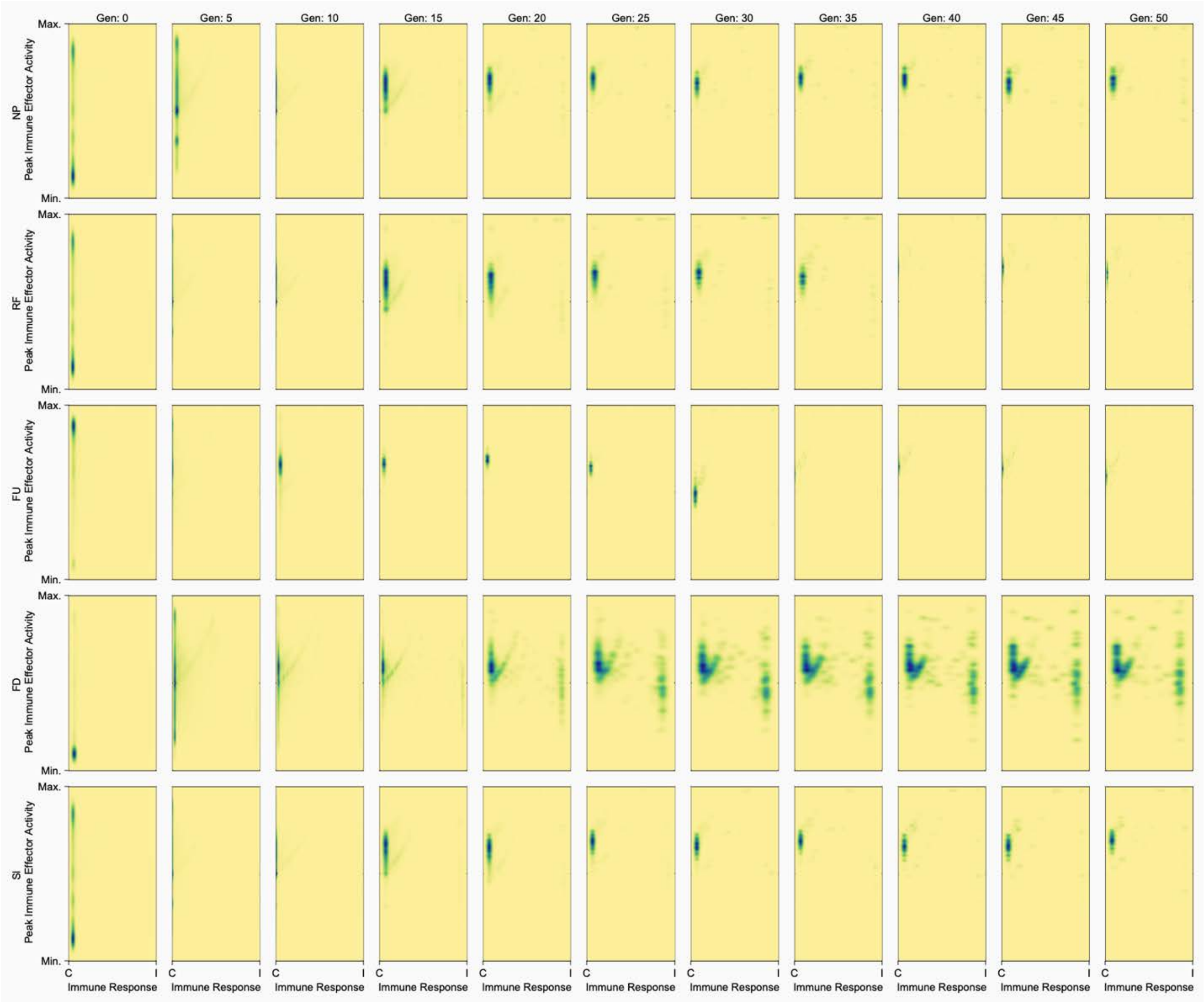


**Supplemental Figure 8c:** Magnitude of immune response by the proportion of response that is induced in the initial 50 generations of an evolutionary simulation where the chance of infection was 90%. Darker colors indicate more common combinations of magnitude of immune responses and proportion of response induced by parasites.

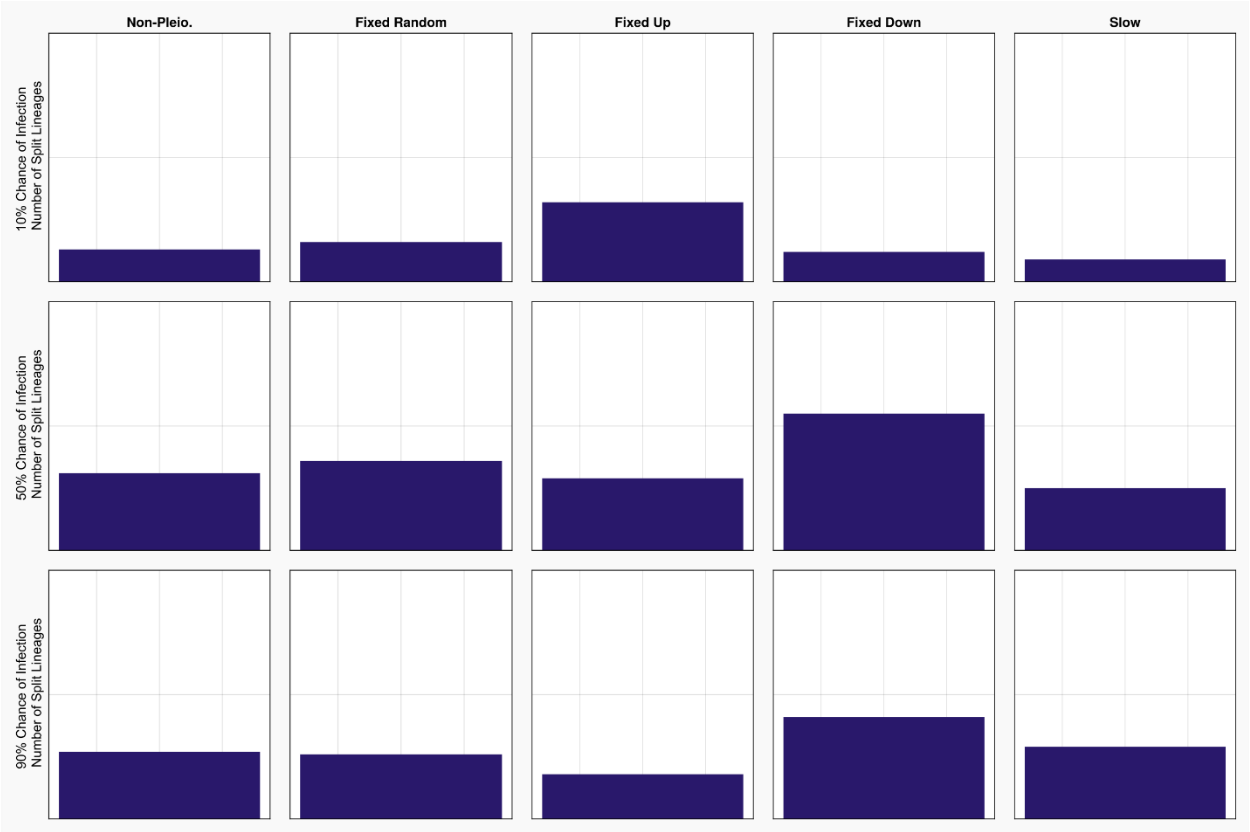


**Supplemental Figure 9:** The proportion of runs where hosts that descended from the same initial host ended up with immune systems with different response dynamics (i.e. a split or bifurcated lineage) in the final generation of the simulation. We refer to hosts that share an ancestor but do not share immune response dynamics as being a part of a split lineage.
